# Human memory CD4^+^ T cells recognize non-infected macrophage bystanders exposed to *Mycobacterium tuberculosis*-infected cells

**DOI:** 10.64898/2026.09.19.752467

**Authors:** Vinicius G. Suzart, Robert Schauner, Scott M. Reba, Daniel P. Gail, Gwendolyn Swarbrick, David M. Lewinsohn, Deborah A. Lewinsohn, Bianca Lepe, Bryan D. Bryson, Samuel M. Behar, Christopher M. Sassetti, Michael L. Freeman, W. Henry Boom, Clifford Harding, Stephen M. Carpenter

**Affiliations:** Division of Infectious Diseases and HIV Medicine, Department of Medicine, Cleveland, OH, USA; Department of Pathology, University Hospitals Cleveland Medical Center, Case Western Reserve University School of Medicine, Cleveland, OH, USA; Portland VA Medical Center, Portland, Oregon, USA; Pulmonary & Critical Care Medicine, Oregon Health & Science University, Portland, Oregon, USA; Molecular Microbiology and Immunology, Oregon Health & Science University, Portland, Oregon, USA; Division of Infectious Diseases, Department of Pediatrics, Oregon Health and Science University, Portland, Oregon, USA; Department of Biological Engineering, Massachusetts Institute of Technology, Cambridge, MA, USA; Ragon Institute of Mass General, Harvard, and MIT, Cambridge, MA, USA; Department of Microbiology, University of Massachusetts Chan Medical School, Worcester, MA, USA

**Keywords:** *Mycobacterium tuberculosis*, TB, memory, CD4, T cell, recognition, infected macrophage, bystander, IFNγ, extracellular vesicles, soluble antigens, type VII secretion system, decoy

## Abstract

Control of *Mycobacterium tuberculosis* (Mtb) infection requires CD4^+^ T cell recognition of infected macrophages. However, T cells also colocalize with non-infected macrophages in granulomas. We investigated whether these bystander macrophages present Mtb antigens and shape human CD4^+^ T cell responses. Using *ex vivo* co-culture systems, non-infected monocyte-derived macrophages (MDMs) were exposed to Mtb-infected MDMs or infection-conditioned supernatants before co-incubation with autologous memory CD4^+^ T cells from individuals with latent Mtb infection (LTBI). Bystander macrophages activated memory CD4^+^ T cells through MHC-II-dependent antigen presentation. Single-cell T cell receptor (TCR) sequencing and TCR-transduced reporter cell lines identified Mtb-specific clonotypes recognizing both infected and bystander macrophages, as well as clonotypes preferentially recognizing infected cells. Strikingly, a subset of TCRs recognized infected but not bystander macrophages. Antigen transfer occurred through soluble Mtb proteins rather than extracellular vesicles. Compared with responses to infected macrophages, bystander macrophages induced attenuated effector responses. These findings reveal antigen-specific recognition of bystander macrophages and suggest that antigens preferentially presented by infected cells may inform TB vaccine design.

## Introduction

In 2024, tuberculosis (TB) was diagnosed in an estimated 10.7 million people, including over 10,000 in the United States, and 1.23 million people died from TB^1^. CD4^+^ T cells are essential to controlling Mtb infection and preventing TB^2–6^. Their central role is highlighted by an up to 20-fold increased risk of TB in people living with HIV, which is greatest in those with low CD4 counts^4,7^. Effector CD4^+^ T cell responses specific for Mtb are first generated in lung-draining lymph nodes (LNs) by non-infected conventional dendritic cells (DCs) that have acquired Mtb antigens from infected cells^8–12^. However, direct recognition of Mtb-infected macrophages by antigen-specific effector CD4^+^ T cells in the lungs is required for control of Mtb infection^9,13–16^. Antigen-specific T cell recognition is mediated through TCR and peptide-major histocompatibility complex class II (pMHC-II) interactions, and TB control is linked to the expression of T cell effector functions including the secretion of GM-CSF, TNF, and IFNγ^3,5,17–19^. However, the role of T cell responses to non-infected macrophages that have acquired Mtb antigens from infected cells remains underexplored, particularly in humans^20,21^.

Pulmonary immune responses to Mtb, including granuloma formation, are dynamic, heterogeneous, and incompletely characterized in humans. Granulomas linked to control of Mtb demonstrate co-localization of T cells and myeloid cells, whereas necrotizing granulomas contain caseous cores harboring numerous bacilli surrounded by epithelioid macrophages and an outer rim of lymphocytes^22–25^. In human lung resection and autopsy specimens, CD4^+^ T cells are frequently observed at the periphery of necrotizing granulomas, adjacent to non-infected macrophages rather than cells containing Mtb^23,25,26^. Low dose Mtb infection in non-human primates (NHPs) shows a architecture, with bacilli concentrated centrally and T cells largely confined to the peripheral regions, adjacent to non-infected macrophages^25–28^. These observations raise the possibility that T cells engage non-infected “bystander” macrophages during TB, particularly when bacterial restriction is poor.

Colocalization of T cells with non-infected macrophages suggests that these bystander cells may present Mtb antigens to T cells. In mice, non-infected DCs can activate Mtb-specific CD4^+^ T cells after acquiring soluble antigens released by infected cells through a kinesin motor complex-dependent mechanism^12,29^. Although antigen transfer to DCs in LNs initiates T cell priming, whether analogous antigen transfer to bystander macrophages elicits and shapes T cell responses in granulomas remains unclear.

In this study, we used *ex vivo* co-culture systems to investigate the extent to which human memory CD4^+^ T cells from healthy individuals with LTBI are activated by non-infected bystander macrophages. Using three different approaches, we found that memory CD4^+^ T cells upregulate activation-induced markers (AIMs) in response to bystander macrophages in a TCR-pMHC-II-dependent manner. Single-cell TCR sequencing of memory CD4^+^ T cells activated in response to bystander macrophages revealed Mtb-specific TCR clonotypes, indicating antigen-specific responses. We then evaluated whether antigen transfer involved extracellular vesicles or soluble proteins, identified Mtb antigens exported from infected cells, and compared the effector programs of T cells responding to infected versus bystander macrophages. We propose that T cells targeting antigens presented preferentially by infected macrophages may represent a particularly protective subset. These results provide a framework for rational TB vaccine design.

## Results

### Human memory CD4^+^ T cells recognize non-infected bystander macrophages exposed to Mtb-infected cells

To investigate CD4^+^ T cell recognition of non-infected macrophages serving as bystanders (BYST) to Mtb-infected cells, we used primary human T cells and autologous MDMs from healthy individuals with LTBI. CD14^+^ cells isolated from PBMCs were differentiated into M1-like MDMs and infected with yellow fluorescent protein (YFP)-expressing Mtb strain H37Rv (YFP-Rv)^30–32^. After 24 h, infected (INF YFP+) and bystander YFP-negative (BYST YFP-) MDMs were separated by flow sorting, re-plated, and subsequently co-cultured with autologous memory (CD45RA^Lo^) CD4^+^ T cells (**Fig. 1A and Supplementary Fig. 1A**). We focused on memory CD4^+^ T cells because this compartment is enriched for Mtb-specific cells in individuals with LTBI^31,33–35^. T cell activation was measured by CD69 and CD40-ligand (CD40L) co-expression^30,31,35–37^ (**Supplementary Fig. 1B**). Memory CD4^+^ T cells responded to INF YFP+ MDMs, as expected, and a substantial fraction also responded to BYST YFP-MDMs (**Fig. 1B and 1C**). Anti-MHC-II mAb blockade reduced activation in response to BYST YFP-MDMs, indicating TCR-pMHC-II-dependent activation (**Fig. 1B and 1C**). To assess residual infection among BYST cells, we measured CFU immediately after sorting and detected ∼1,000-fold fewer Mtb CFU among BYST YFP-MDMs despite only ∼2-fold fewer activated T cells (**Fig. 1C and 1D**). Thus, memory CD4^+^ T cells recognize non-infected bystander macrophages exposed to Mtb-infected cells in a TCR-pMHC-II-dependent manner.

**Figure 1.**
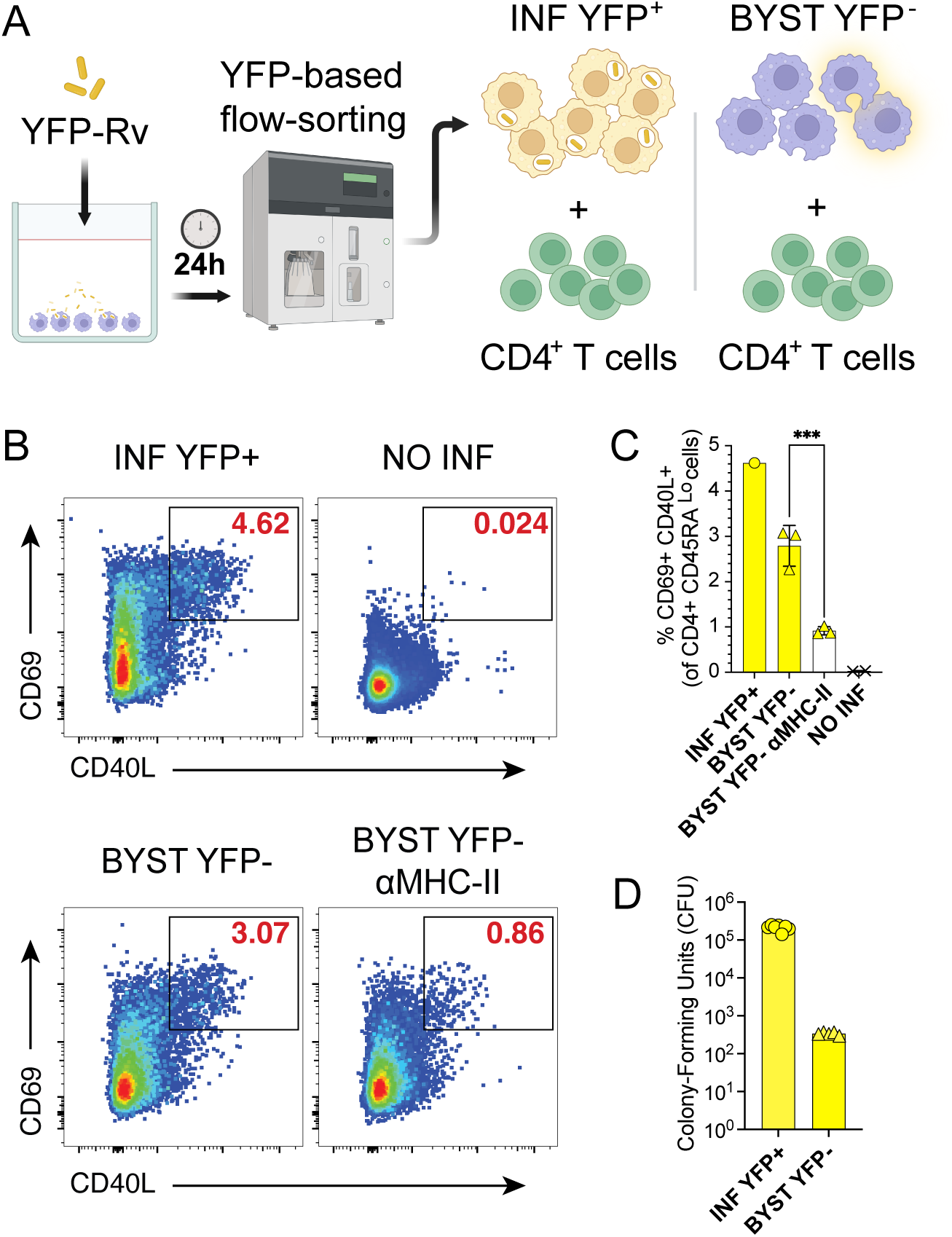
Human memory CD4^+^ T cells recognize non-infected bystander macrophages exposed to Mtb-infected cells. **(A)** Graphical representation of YFP-Mtb macrophage flow sorting in tandem with T cell co-culture assays. Created with Biorender. **(B)** Representative flow cytometry plots and **(C)** summary bar graph showing the proportion (mean ± s.d.) of memory (CD45RA^Lo^) CD4^+^ T cells that co-express CD69 and CD40L after 16-18 h in co-culture with non-infected (NO INF), infected (INF YFP+), or bystander (BYST YFP-) macrophages, ± αMHC-II mAb blockade. Dots indicate replicates from a representative of 3 independent experiments (3 individual participants) containing 1-3 replicates per condition. One-way ANOVA with Sidak’s post-test corrected for multiple comparisons was used to determine statistical significance. (**D)** Summary bar graphs of colony-forming units (CFU) from INF YFP+ or BYST YFP-macrophages 24 h post-infection immediately after sorting. * p<0.05; ** p < 0.01; *** p < 0.001; **** p < 0.0001.

### Human CD4^+^ T cells recognize bystander macrophages in the absence of direct contact with Mtb-infected cells

Bystander macrophages could acquire Mtb antigens through several mechanisms, including direct contact with infected macrophages, efferocytosis, extracellular vesicles (EVs), soluble antigens released into supernatants, or exposure to the Mtb inoculum^12,21,38,39^. To determine whether direct contact with infected cells is required, we exposed BYST macrophages only to conditioned supernatants from INF cells using a transwell system that physically separates infected and bystander macrophages while allowing soluble mediators to pass, modeling the spacial organization of inner core (infected) and outer rim (non-infected) macrophages in granulomas (**Fig. 2A**). Upper chambers containing MDMs plated on 0.4 μm-pore membranes were added to wells containing INF macrophages after the Mtb inoculum was washed out and were incubated for 24 h before addition of CD4^+^ T cells. Memory CD4^+^ T cells responded to both INF and BYST macrophages (**Fig. 2B and 2C**), and αMHC-II blockade significantly reduced responses to BYST macrophages, indicating TCR-pMHC-II-dependent activation (**Fig. 2B and 2C**). BYST macrophages in upper transwell chambers were rarely YFP+ after exposure to YFP-Rv-infected macrophages and contained ∼10,000-fold fewer CFU than INF macrophages (**Fig. 2D and 2E**). These results demonstrate that memory CD4^+^ T cell responses to bystander macrophages occur without direct exposure to infected cells, or the Mtb inoculum, and are consistent with antigen-specific recognition.

**Figure 2.**
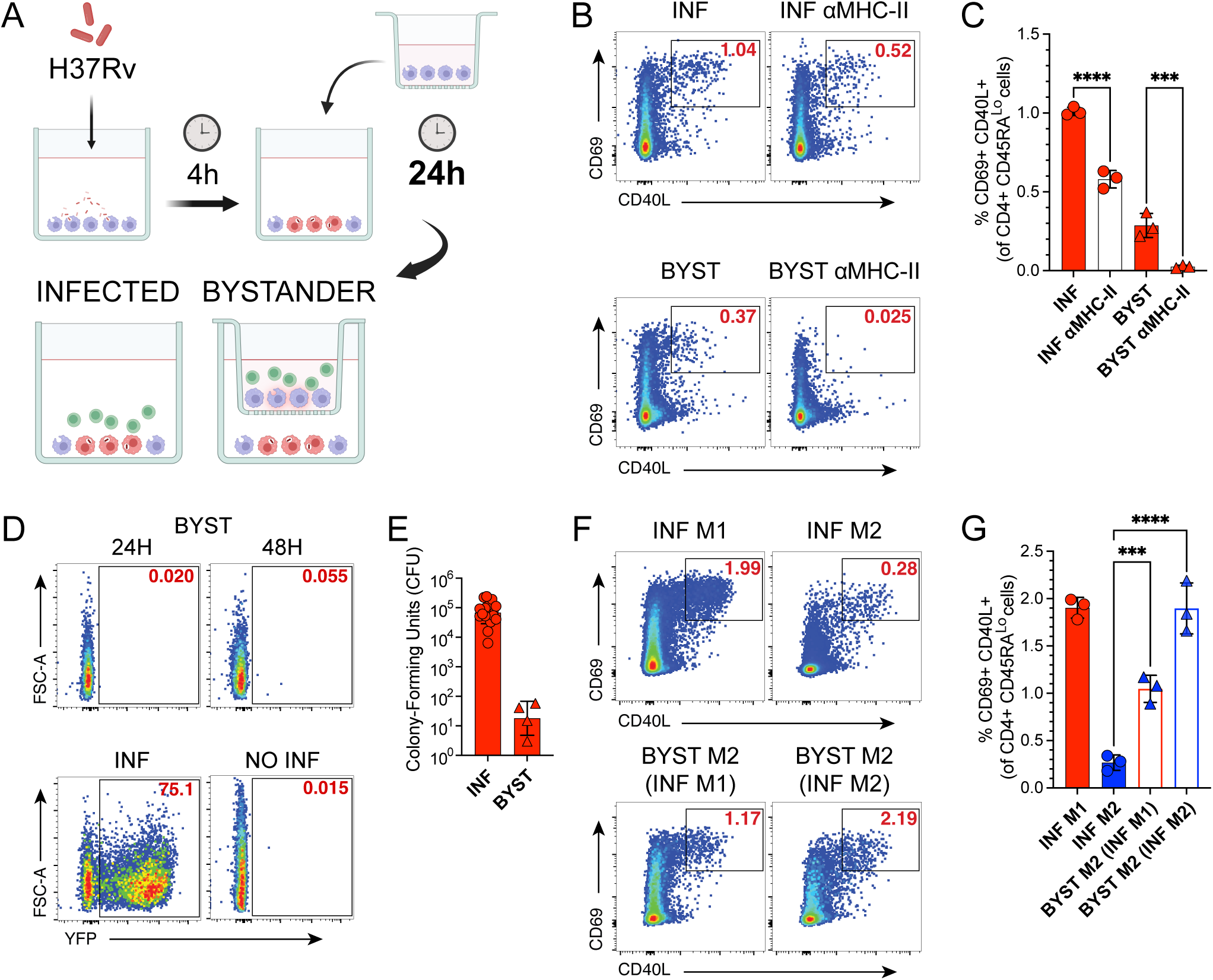
Human CD4^+^ T cells recognize bystander macrophages in the absence of direct contact with Mtb-infected cells. **(A)** Graphical representation of macrophage infection and T cell co-culture. Created with Biorender. **(B)** Representative flow cytometry plots and (**C)** summary bar graph of the proportion (mean ± s.d.) of memory (CD45RA^Lo^) CD4^+^ T cells co-expressing CD69 and CD40L after 16-18 h in co-culture with INF or BYST macrophages ± αMHC-II mAb blockade. **(D)** Representative flow cytometry plots of YFP expression by non-infected (NO INF), YFP-Rv-infected (INF) or BYST macrophages 24-48 h after YFP-Rv infection. **(E)** Summary bar graph of colony-forming units (CFU) (mean ± s.d.) from INF M1 or BYST M1 macrophages from transwell system 24 h post-infection. **(F)** Representative flow cytometry plots and **(G)** summary bar graph of proportion (mean ± s.d.) of memory (CD45RA^Lo^) CD4^+^ T cells that co-express CD69 and CD40L after 16-18 h in co-culture with INF M1- or M2-like macrophages, or with BYST M2-like macrophages exposed to either INF M1-or M2-like macrophages. Dots indicate replicates from a representative of 3-9 independent experiments (3-5 individual participants). One-way ANOVA with Sidak’s post-test corrected for multiple comparisons was used to determine statistical significance. * p<0.05; ** p < 0.01; *** p < 0.001; **** p < 0.0001.

We previously showed that M1- and M2-like human MDMs differ in their ability to activate CD4^+^ T cells after Mtb infection, although both present exogenous peptides, H37Rv whole-cell lysate, or gamma-irradiated Mtb^31,40^. We therefore asked whether bystander M2-like MDMs could activate CD4^+^ T cells despite poor activation when directly infected. In the transwell system, BYST M2-like MDMs were exposed to INF M1- or M2-like MDMs and then compared across conditions. As expected, INF M2-like MDMs were poorly recognized compared with INF M1-like MDMs (**Fig. 2F and 2G**). However, M2-like bystander macrophages elicited robust CD4^+^ T cell responses after exposure to either INF M1- or INF M2-like macrophages. Thus, Mtb-infected M2-like MDMs can transfer antigens to bystander MDMs despite their limited ability to directly activate CD4^+^ T cells.

### CD4^+^ T cell recognition of bystander macrophages is Mtb antigen-specific

We recently reported the antigen specificities of TCRs sequenced from human memory CD4^+^ T cells activated by Mtb-infected macrophages from individuals with LTBI^35^. To test whether Mtb-specific clonotypes also respond to bystander macrophages, we flow-sorted activated memory CD4^+^ T cells responding to BYST or INF autologous MDMs in the transwell system and performed single-cell TCR sequencing (**Fig. 2A and Fig. 3A-D**). These TCRs were combined with sequences from our prior study and clustered by CDR3β similarity using grouping of lymphocyte interactions by paratope hotspots V2 (GLIPH2)^35–37,41^. After removing GLIPH2 groups linked to non-Mtb specificities or control peptide responses (**Extended Data 1**), 246 groups containing 646 unique TCRs met statistical criteria and represented TCRs from ≥3 participants. Surprisingly, 60% of these groups contained clonotypes responding to both INF and BYST macrophages (**Fig. 3B and 3C**). We also identified TCRs responding to the Mtb peptide megapool (MTB300)^34^ or Mtb lysate, but not to infected macrophages (**Fig. 3A**), half of which (6% of TCRs) responded to bystander macrophages only, whereas 29% of clonotypes were linked to INF but not BYST macrophages (**Fig. 3B and 3C**). Strikingly, several GLIPH2 groups linked to BYST and INF macrophage responses contained TCRs we previously identified as Mtb-specific (blue font on the x-axis), from which we had already cloned representative TCRs from GLIPH2 groups linked to responses to Mtb peptides or infected macrophages^35^. These included the S%SGTKYNE (PE18/PE19_1-15_-specific) GLIPH2 group, SPGTESN%P and S%GTESNQP (EspA_301-315_), SSPGQGG%NYG (EspC_74-88_), and %MPE (EsxG/EsxS_49-63_), and previously identified GLIPH2 groups for yet-to-be defined antigens (SFVDS%YE, SES%GGSNQP, S%ADSNQP, SRGTGG%YE), where “%” represents any amino acid (**Fig. 3D**). We also identified Mce3A_111-125_ (Rv1966) specificity within the SA%RDQP group, which responded to BYST and INF macrophages, by screening overlapping peptide pools from 68 Mtb antigens in the IMPAC-TB library (**Supplementary Fig. 2A-C and Extended Data 2**). In contrast, certain EsxB_52-66_-specific TCRs (R%SGGEAK%NI and RA%GGEAKNI groups, blue font on the x-axis) responded only to Mtb-infected macrophages^35,41^ (**Fig. 3D**). These results indicate that responses to non-infected bystander macrophages include Mtb-specific CD4^+^ T cell clonotypes.

**Figure 3.**
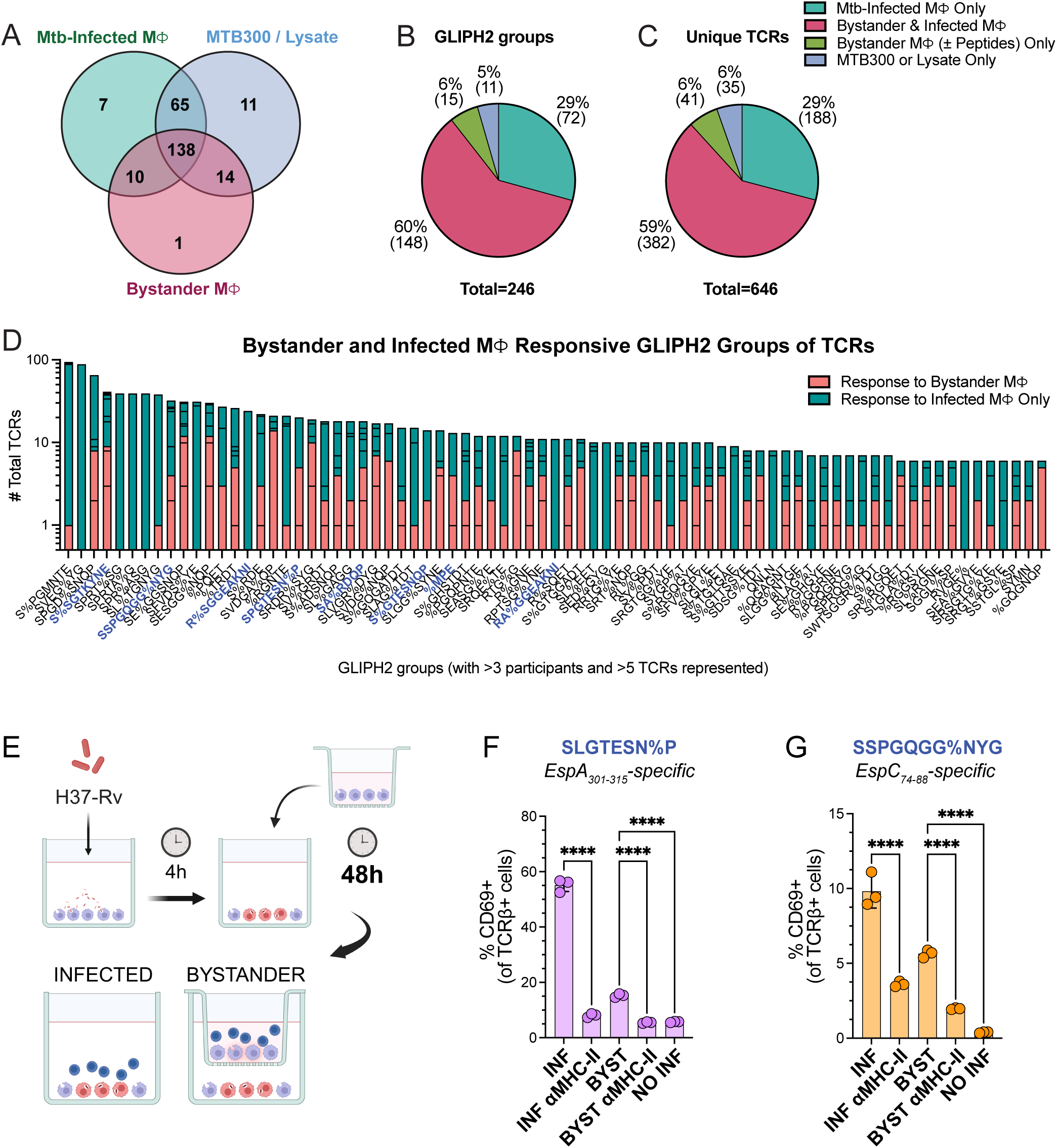
CD4^+^ T cell recognition of bystander macrophages is Mtb antigen-specific. **(A)** Venn diagram indicating the number of GLIPH2 groups containing TCRs linked to responses to Mtb-infected (green), bystander macrophages (pink), or after addition of MTB300/Lysate (blue). **(B, C)** Pie charts indicating the percentage (and number) of GLIPH2 groups **(B)** and unique TCRβ sequences **(C)** linked to responses to Mtb-infected (teal), bystander (green), bystander and infected (pink), or after addition of MTB300/Lysate (blue). **(D)** Bar graphs indicating the number of GLIPH2 groups containing TCRs from ≥ 3 participants) linked to responses to bystander (orange) or to Mtb-infected macrophages only (green). GLIPH2 groups containing TCRs with confirmed specificity to Mtb antigens are in blue text on x-axis. Data combined from 5 individual participants in 2 independent experiments and aggregated with previously published TCRs^35–37,41^ (n=116 unique participants) for GLIPH2 analysis. **(E)** Graphical representation of macrophage infection and Mtb-specific SKW-3 cell lines co-culture. Created with Biorender. **(F, G)** Summary bar graphs of the proportion (mean ± s.d.) of **(F)** EspA_301-315_- and **(G)** EspC_74-88_-specific TCR-transduced SKW-3 cells, respectively, expressing CD69 16-18 h after co-culture in upper transwell chambers that were exposed to infected macrophages for 48 h ± αMHC-II mAb blockade. GLIPH2 groups containing the respective Mtb-specific TCRs are in blue text. One-way ANOVA with Sidak’s post-test corrected for multiple comparisons was used to determine statistical significance. * p<0.05; ** p < 0.01; *** p < 0.001; **** p < 0.0001.

We next sought to validate antigen-specific recognition of bystander macrophages using TCR-transduced SKW-3 cell lines which expresses CD69 upon TCR engagement^35^. Since we detected EspA_301-315_-specific (SPGTESN%P and S%GTESNQP) and EspC_74-88_-specific (SSPGQGG%NYG) TCRs among the responses to BYST MDMs in upper chambers, we tested SKW-3 cell lines expressing those respective TCRs for recognition of BYST MDMs by assessing activation after co-culture with BYST MDMs in the upper chambers of the transwell system exposed to INF MDMs for 48 h (**Fig. 3E**). EspA_301-315_-specific cells were activated by BYST macrophages at reduced but detectable levels compared with INF macrophages, and activation was prevented by αMHC-II blockade (**Fig. 3F**). EspC_74-88_-specific cells were similarly activated by BYST macrophages in an MHC-II-dependent manner (**Fig. 3G**). Activation of EspC_74-88_-specific cells increased with longer BYST exposure to INF macrophages, peaking at 48 h, consistent with antigen accumulation by bystander macrophages (**Supplementary Fig. 2D**). Together, these data confirm that Mtb antigen-specific CD4^+^ T cells recognize non-infected bystander macrophages.

### Recognition of bystander macrophages by Mtb-specific CD4^+^ T cells occurs through the transfer of soluble Mtb antigens from infected cells

We next sought to determine how bystander macrophages acquire Mtb antigens in the absence of direct contact with infected cells. EVs, including exosomes, microvesicles, apoptotic bodies, and bacterial vesicles (BVs), could transfer Mtb antigens due to their prominent role in intercellular communication and have been implicated in intercellular communication during mycobacterial infection^42–46^. Due to the 0.4 μm pore size in the filter separating the transwell chambers, the average diameters of BVs (0.06 to 0.3 μm) and host-derived exosomes (0.03 to 0.15 μm), microvesicles (0.1 to 1 μm), and apoptotic bodies (1 to 5 μm) suggested the transwell filter would permit BVs and exosomes but limit larger EVs. We measured the presence of exosomes in supernatants conditioned by Mtb-infected macrophages after 0.22 μm-filtration using surface receptors commonly expressed on exosomes, the tetraspanins CD9, CD63, and CD81, in addition to MHC-II (HLA-DR/DP/DQ) (**Fig. 4A and Supplementary Fig. 3**). After 24-48 h, exosomes were more abundant in supernatants from M1-than M2-like MDMs, irrespective of Mtb infection (**Fig. 4A**), making exosomes candidate mediators of antigen transfer from infected M1-like MDMs.

**Figure 4.**
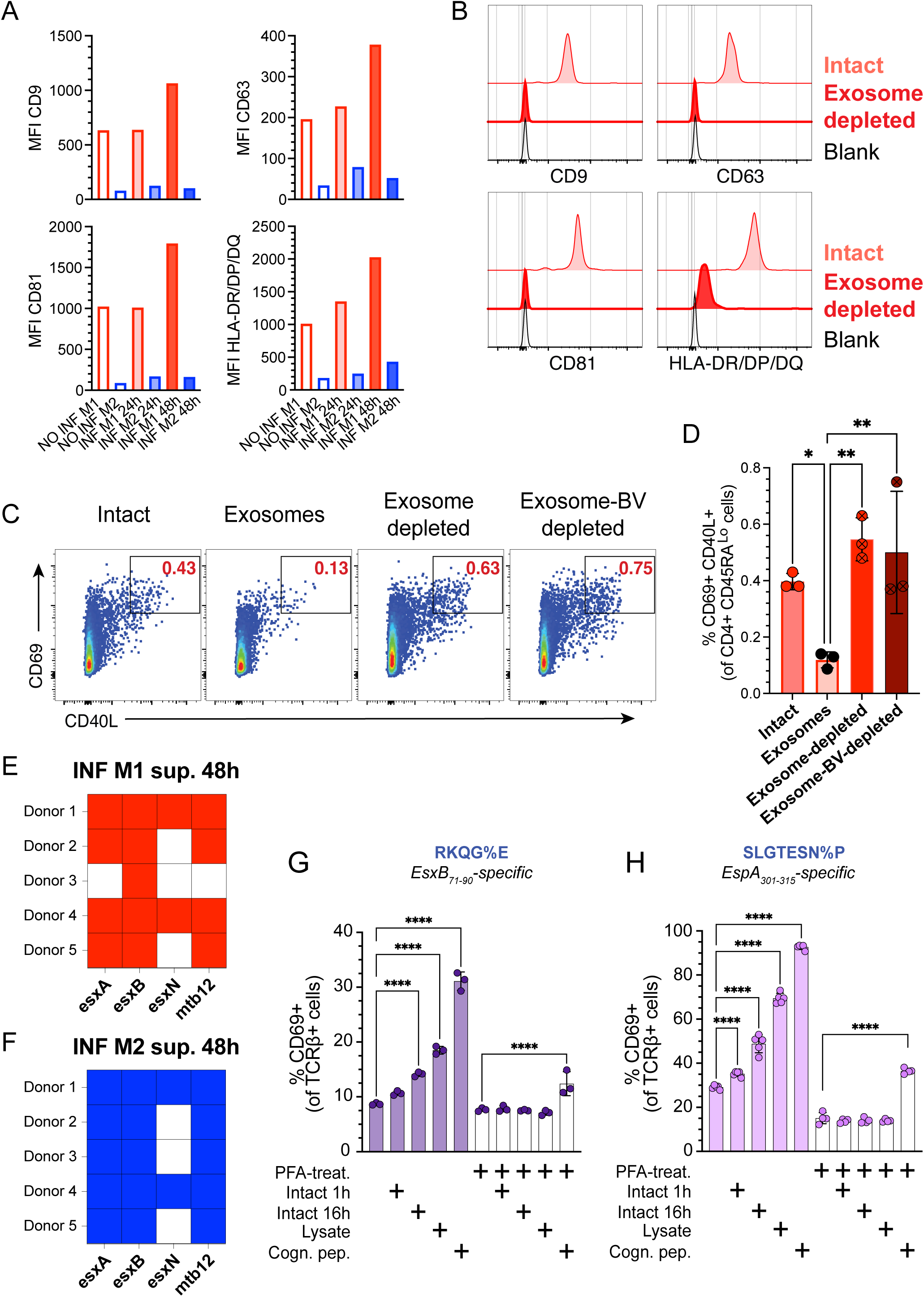
Recognition of bystander macrophages by Mtb-specific CD4^+^ T cells occurs through the transfer of soluble Mtb antigens from infected cells. **(A)** Summary bar graph of expression of exosome surface tetraspanin receptors CD9, CD63, and CD81, and MHC-II (HLA-DRDPDQ) (individual values) from 0.22 μm-filtered serum-free supernatants conditioned by INF M1- or M2-like macrophages 24 or 48 h post-infection. **(B)** Representative histograms showing median fluorescence intensity (MFI) of exosome surface tetraspanin receptors CD9, CD63, and CD81, and MHC-II (HLA-DR/DP/DQ) of 0.22 μm-filtered (intact) or exosome-depleted supernatants. (**C)** Representative flow cytometry plots and **(D)** summary bar graph of the proportion (mean ± s.d.) of memory (CD45RA^Lo^) CD4^+^ T cells co-expressing CD69 and CD40L AIMs after co-culture for 18 h with non-infected M1-like MDMs treated with either 0.22 μm-filtered conditioned supernatant (intact), exosome, exosome-depleted, or exosome-BV-depleted supernatant fractions for 24 h prior to T cell-macrophage co-culture. Data are from a representative of 2 independent experiments (2 individual participants). **(E, F)** Summary plot of Mtb proteins identified by mass spectrometry in 0.22 μm-filtered conditioned supernatant of INF M1- or M2-like MDMs 48 h post-infection. **(G, H)** Summary bar graph of the proportion (mean ± s.d.) of **(G)** EsxB_71-90_- and **(H)** EspA_301-315_-specific TCR-transduced SKW-3 cells expressing CD69, 16-18 h after co-culture with MDMs ± 0.25% PFA-fixation followed by treatment with either Mtb lysate, cognate EsxB_71-90_ or EspA_301-315_ peptide, or intact conditioned supernatant harvested 48 h post infection. Data are from a representative of 2 independent experiments. One-way ANOVA with Sidak’s post-test corrected for multiple comparisons was used to determine statistical significance. * p<0.05; ** p < 0.01; *** p < 0.001; **** p < 0.0001.

To test whether EVs mediate antigen transfer, we treated non-infected macrophages with supernatants conditioned by Mtb-infected M1-like MDMs before or after EV depletion. Exosomes were depleted from 0.22 μm-filtered conditioned supernatants by immunomagnetic selection using anti-CD9, anti-CD63, and anti-CD81 microbeads, and depletion was confirmed by loss of HLA-DR and tetraspanin receptor signal (**Fig. 4B**). Remaining small microvesicles and BVs were then depleted by ultracentrifugation. After treatment with intact, exosome-depleted, or total EV-depleted conditioned supernatants, non-infected macrophages activated similar numbers of memory CD4^+^ T cells, whereas purified exosomes induced little to no activation (**Fig. 4C and 4D**). Together, these data indicate that recognition of bystander macrophages occurs through transfer of soluble Mtb antigens from infected macrophages and does not require EVs.

To define the breadth of antigens released by infected macrophages, we performed mass spectrometry on supernatants conditioned by infected M1- and M2-like MDMs 48 h post-infection. We identified Mtb proteins in these conditioned supernatants, all cell wall or secreted proteins including type VII secretion system (T7SS) substrates [EsxA (ESAT-6, Rv3875), EsxB (CFP-10, Rv3874), EsxN (Rv1793)], and the general secretory pathway (Sec) substrate mtb12 (cfp2, Rv2376c) (**Fig. 4E and 4F and Extended Data 3**). These data show that infected MDMs release Mtb antigens into conditioned supernatants encountered by bystander macrophages, supporting antigen acquisition and presentation by non-infected bystander cells.

To determine whether Mtb antigens were transferred in the form of whole proteins or processed peptides, we treated non-infected macrophages with or without light paraformaldehyde (PFA) fixation and exposed them for 1 h or 16 h to conditioned supernatants from infected macrophages^47^. Since mass spectrometry detected EsxB in multiple donor conditioned supernatants, we tested the EsxB_71-90_-specific (RKQG%E) and EspA_301-315_-specific (SPGTESN%P and S%GTESNQP) TCR-transduced cell lines with macrophages treated with conditioned supernatants, lysate, or cognate peptide. The EspA-specific TCR-transduced cell line was then co-cultured with conditioned supernatant-treated macrophages and activation was assessed. We reasoned that PFA-fixed non-infected MDMs would activate TCR-transduced cells only if cognate antigens were already present as processed peptides available for direct MHC-II loading, especially for the 1 h exposure. We found that the EsxB_71-90_- and EspA_301-315_-specific cells responded to unfixed MDMs treated with conditioned supernatants, lysate, or cognate peptide, but not to PFA-fixed MDMs treated with conditioned supernatants unless cognate peptide was added (**Fig. 4G**). Together with the greater T cell activation after 16 h vs. 1 h, these findings indicate that non-infected macrophages take up and process soluble EsxB and EspA proteins from conditioned supernatants, supporting transfer of unprocessed Mtb proteins from infected to non-infected human macrophages.

### T cell clusters enriched for Mtb-specific TCRs express a multifunctional effector program

We next compared effector phenotypes of memory CD4^+^ T cells activated by INF or BYST macrophages using single-cell transcriptomics (scRNAseq) of flow-sorted cells. The integrated dataset included 41,190 high-quality CD4^+^ T cells. Among genes differentially expressed after INF recognition, *IFNG* and *IL22* were significantly upregulated (**Fig. 5A and Supplementary Fig. 4**). Louvain clustering identified 12 clusters visualized by UMAP (**Fig. 5B and Extended Data 4**). Mapping αβTCR sequences to cell barcodes showed that clusters 0 and 4 were enriched for clonally expanded CD4^+^ T cells responding to INF or BYST macrophages (**Fig. 5C**). Cluster 4 preferentially expressed *IFNG, IL2, CSF2* (GM-CSF), *TNF, GZMB,* and *CCL20*, consistent with the multifunctional effector CD4^+^ T cell program we recently reported for Mtb-specific clonotypes^35^. Cluster 0 cells preferentially expressed *CXCR4, DPP4, SELL, RGS1* and *CCR6,* indicating features of antigen-experienced Th17/Th1* cells^33,48–50^ (**Fig. 5D and Extended Data 4**). Clusters 0 and 4 were both populated by a greater proportion of T cells responding to INF macrophages (**Fig. 5E**). Cluster 7 was enriched for BYST-responsive cells but largely contained non-expanded TCRs, suggesting non-specific activation by this TNF-producing subset. Finally, we mapped TCRs from GLIPH2 groups responsive to Mtb or “control” peptides, using prior annotations, including those from the immune epitope database (iedb.org)^35–37,41,51^. While both clusters 0 and 4 contained antigen-specific clonotypes, the Mtb-specific TCRs preferentially populated cluster 4, particularly among the responses to infected macrophages (**Fig. 5F**). These results indicate that Mtb-specific CD4^+^ T cells responding to infected or bystander macrophages preferentially co-express *IFNG, IL2, TNF, and CSF2*, and is enriched among responses to infected cells.

**Figure 5.**
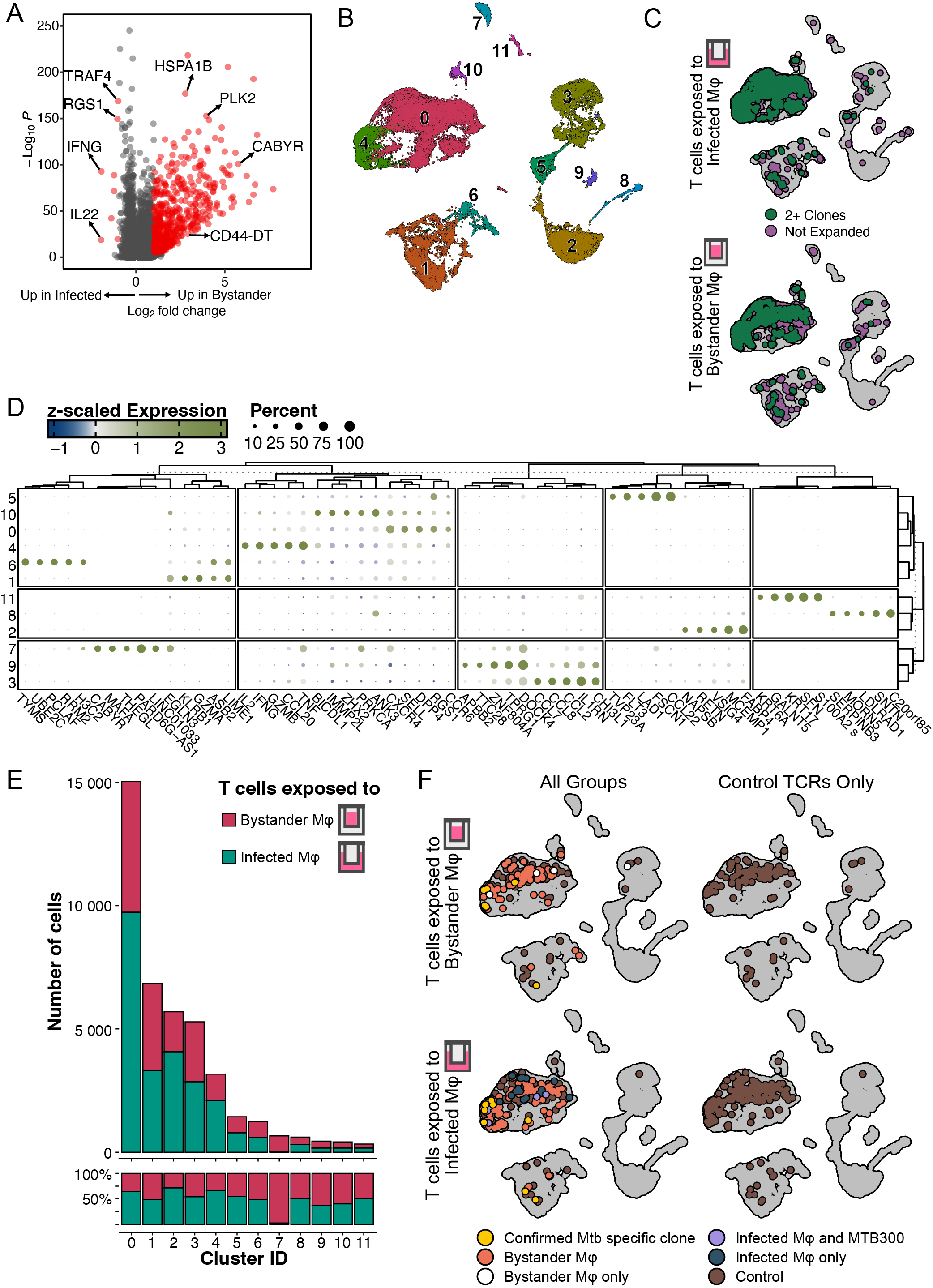
T cell clusters enriched for Mtb-specific TCRs express a multifunctional effector program. **(A)** Volcano plot comparing log_2_ fold change of genes upregulated in CD69^+^ CD40L^+^ T cells co-cultured with INF or BYST macrophages in our transwell system for 16-18 h. Red dots represent genes with significant upregulation in each condition using MAST (adjusted p-value < 0.05 and abs(log_2_ fold change) > 0.5)). **(B)** UMAP plot showing Louvain clustering of the flow-sorted activated CD4^+^ T cell populations in response to INF or BYST macrophages after quality control and integration, combined from 5 individual participants in 2 independent experiments. **(C)** Split UMAP plots for each condition mapping all cells with sequenced TCR clonotype present in one (purple) or 2 or more cells (green). **(D)** Dot plot of the top 10 upregulated genes per cluster with adjusted p-value < 0.05 and >10% expression in a given cluster. **(E)** Bar graph indicating the number (top) or proportion (bottom) of T cells obtained from co-culture with INF (green) or BYST (pink) macrophages populating each cluster and respective relative percentages. **(F)** Split UMAP plots showing cells from BYST (top) and INF (bottom) highlighting GLIPH2 groups containing TCRs identified in response to INF (green), INF and MTB300-treated (purple), BYST (orange), BYST only (white), confirmed Mtb-specific (yellow), and GLIPH2 groups containing TCRs previously linked to responses to control (viral or vaccine) peptide responses^35^ (brown, left and right).

### Fewer CD4^+^ T cells secrete GM-CSF and IFNγ in response to bystander macrophages

Since clusters 0 and 4 mapped the majority of clonally-expanded TCRs, and cluster 7 contained greater numbers of T cells responding to BYST macrophages, we sought to compare the expression of effector genes among T cells from each condition in clusters 0, 4, and 7. Although cluster 4 was composed of responses to both BYST and INF macrophages, fewer T cells responding to bystander macrophages expressed the canonical Th1 cytokine, *IFNG*, and transcription factor, *TBX21* (**Fig. 6A**). Similarly, fewer T cells expressed *IL2* and *GZMB*, but not *TNF*, when compared to those responding to infected macrophages. These data suggest that recognition of bystander macrophages is less likely to lead to effector cytokine production.

**Figure 6.**
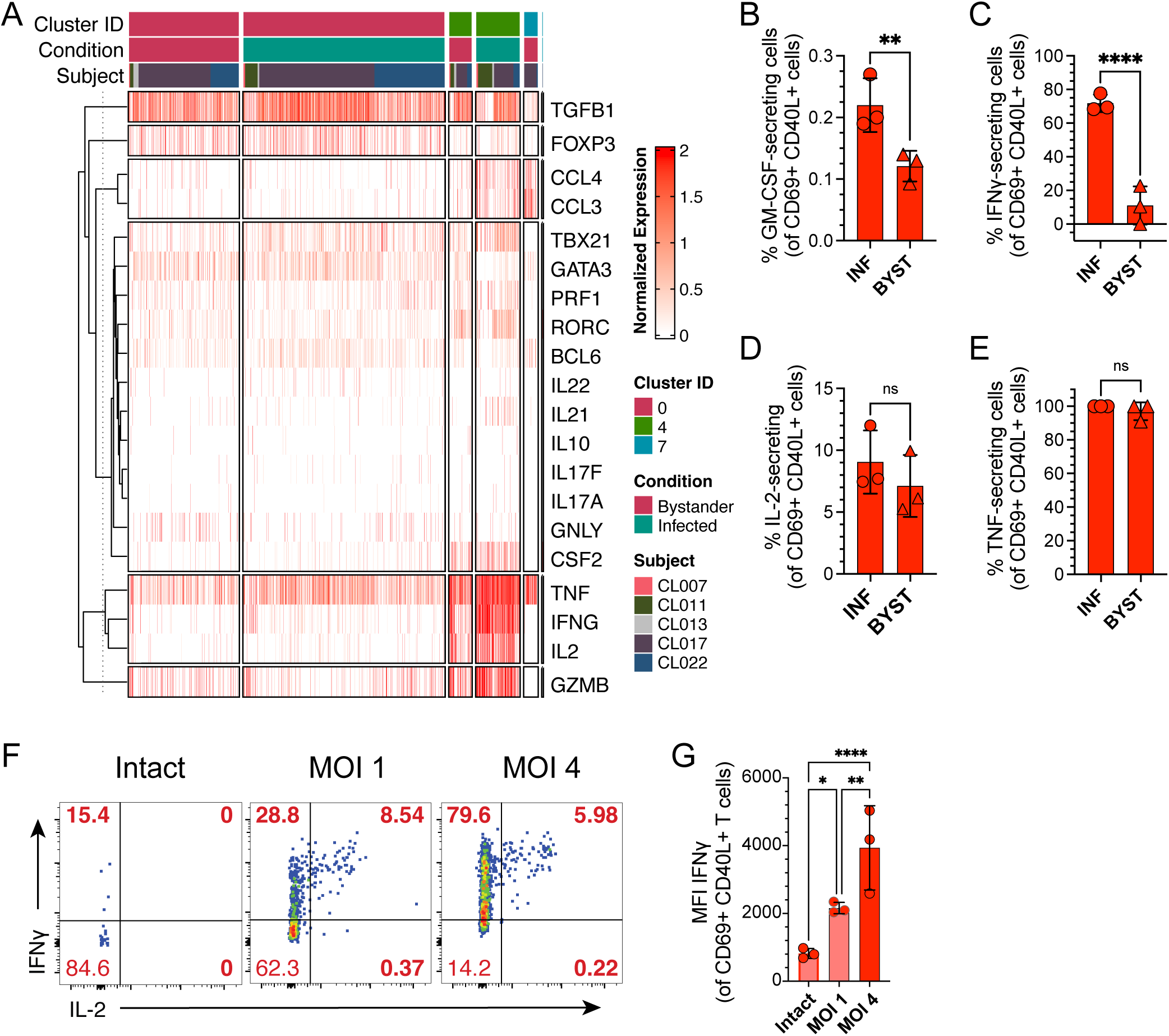
Fewer CD4^+^ T cells secrete GM-CSF and IFNγ in response to bystander macrophages. **(A)** Heatmap representing log-normalized counts of effector genes for T cells obtained from co-culture with INF (green) or BYST (pink) macrophages from clusters 0 (pink), 4 (green) and 7 (blue), respectively. Representative bar graph of proportion (mean ± s.d.) of (**B**) GM-CSF-, (**C**) IFNγ-, (**D**) IL-2-, and (**E**) TNF-secreting T cells, gated on CD69^+^ CD40L^+^ memory (CD45RA^Lo^) CD4^+^ T cells co-cultured with INF or BYST macrophages for 16-18 h from 3 independent experiments (3 individual participants). **(F)** Representative flow cytometry plots of CD69^+^CD40L^+^ memory CD4^+^ T cells secreting IFNγ and IL-2, and **(G)** summary bar graphs of IFNγ MFI (mean ± s.d.), 16-18 h after co-culture with autologous M1-like MDMs treated with infection-conditioned supernatants or Mtb-infected (MOI 1 or 4). Data are from a representative of 6 independent experiments (3 individual participants). One-way ANOVA with Sidak’s post-test corrected for multiple comparisons was used to determine statistical significance. * p<0.05; ** p < 0.01; *** p < 0.001; **** p < 0.0001.

We next sought to compare the capacity of memory CD4^+^ T cells to secrete effector cytokines in response to BYST or INF macrophages at the protein level using a flow cytometry-based cytokine secretion assay. While high TNF secretion and modest IL-2 secretion were similar between conditions, fewer memory CD4^+^ T cells secreted GM-CSF or IFNγ in response to BYST macrophages compared to INF macrophages, recapitulating the results from single-cell transcriptomics (**Fig. 6B-E**). We also detected fewer IFNγ-secreting memory CD4^+^ T cells in response to non-infected MDMs treated with conditioned supernatants compared to macrophages infected at MOIs 1 or 4 (**Fig. 6F and 6G**). Together, our findings indicate that memory CD4^+^ T cell responses to non-infected bystander macrophages express a distinct and attenuated effector profile compared to those responding to Mtb-infected cells.

## Discussion

In this study, we show that human memory CD4^+^ T cells from individuals with LTBI recognize non-infected macrophages exposed to Mtb-infected cells. Using flow sorting of infected and non-infected bystander macrophages, a transwell system to segregate infected from bystander APCs, and the transfer of filtered supernatants conditioned by infected macrophages, we discovered TCR-pMHC-II-dependent CD4^+^ T cell activation in response to bystander cells. Using single-cell TCR sequencing, we identified Mtb-specific TCRs expressed by T cells activated in response to bystander macrophages and confirmed the bystander recognition was antigen specific using TCR-transduced SKW-3 cell lines. Importantly, we captured distinct subsets of TCRs linked to exclusive recognition of Mtb-infected macrophages. EV depletion experiments, proteomic analysis of supernatants conditioned by infected macrophages, and light PFA fixation assays indicate that bystander macrophages acquire soluble, unprocessed Mtb proteins released by infected cells and process them for presentation. Finally, single-cell transcriptomics and cytokine secretion assays showed that T cells responding to bystander macrophages have attenuated effector profiles, including reduced IFNγ production. Together, these findings establish antigen-specific recognition of non-infected bystander macrophages through intercellular transfer of soluble Mtb proteins and suggest that this interaction alters CD4^+^ T cell effector function.

Effective control of Mtb infection was shown to require direct recognition of infected macrophages via TCR-pMHC-II interaction^9,13,14^. Our data show that Mtb-specific CD4^+^ T cells also recognize non-infected bystander macrophages, but express attenuated effector responses characterized by reduced IFNγ production. Prior evidence of suboptimal CD4^+^ T cell activation in Mtb-infected murine lungs^15,16,32^, together with our observation of inefficient recognition of human macrophages expressing an M2-like phenotype upon Mtb infection^31,40^, these findings raise the possibility that antigen presentation by bystander macrophages contributes to limiting CD4^+^ T cell effector function during TB. One possibility is that Mtb-specific memory CD4^+^ T cells encountering antigen on bystander macrophages with insufficient costimulatory receptor expression fail to mount robust effector responses upon subsequent encounter with infected cells, thereby limiting local antimicrobial immunity^52^. Alternatively, bystander macrophages near infected cells could enhance local immunity if they present antigen with appropriate costimulation and signal 3 cytokines. Differences in pMHC-II density, costimulatory molecules, and local cytokines are therefore likely to shape whether antigen transfer promotes protection or pathogenesis. Therefore, intercellular transfer of Mtb antigens to non-infected bystander macrophages in the lungs could either play an important role in pathogenesis of the infection, by misdirecting antigen-specific T cells and limiting their effector responses, or a protective role, by increasing the number of activated CD4^+^ T cells localized to the site of infection. In the “decoy” scenario^14^, antigen presentation by bystander macrophages could disperse or dampen protective T cell responses, facilitating Mtb persistence within granulomas. Determining whether this occurs *in vivo* will require models that recapitulate human granuloma architecture.

Reduced effector responses to bystander macrophages could reflect a tolerant or dysfunctional state such as anergy, which has been linked to the direct effects of lipoarabinomannan on T cells^53^. T cell anergy is characterized by diminished cytokine production in response to antigen exposure^54,55^ and has been described during TB^53,56^. Regulatory cytokines, including IL-10 and TGF-β, are known modulators of T cell function during Mtb infection^57–59^. TGF-β was shown to suppress IFNγ secretion by Mtb-specific CD4^+^ T cells in granulomas^58^. Although we did not directly test TGF-β secretion by bystander macrophages, gene set enrichment analysis of DEGs in activated T cells did not identify enrichment of TGF-β or IL-10 signaling pathways. Instead, T cells responding to bystander macrophages showed reduced *TBX21* expression, which may contribute to diminished IFNγ production. These effects could result from antigen recognition on bystander macrophages with limited costimulation or from exposure to soluble mycobacterial lipids such as lipoarabinomannan^53^. Overall, reduced pro-inflammatory cytokine production by CD4^+^ T cells responding to bystander macrophages is unlikely to reflect lineage diversion and may instead arise from reduced effector Th1 polarization.

Mass spectrometry identified secreted Mtb proteins in supernatants conditioned by infected macrophages, including T7SS and Sec substrates. EspA was not detected in conditioned supernatants, likely due to insufficient sensitivity in the setting of low antigen abundance. However, EspA-specific TCRs were among the most abundant clonotypes responding to infected and bystander macrophages, and EspA_301-315_-specific cells recognized bystander MDMs and unfixed MDMs treated with conditioned supernatants. Thus, failure to detect EspA likely reflects limited analytical sensitivity rather than absence of antigen. Because CD4^+^ T cell activation by bystander macrophages occurred despite physical separation from infected cells, and after EV depletion, our data support a model in which soluble Mtb proteins are transferred and processed by non-infected macrophages. This model is consistent with reports of secreted T7SS peptides eluted from pMHC molecules on infected human APCs^60,61^. Most Mtb-specific TCR clonotypes recognized both infected and bystander macrophages, suggesting that shared pMHC-II complexes can be presented by both cell states. However, some TCRs preferentially or exclusively recognized infected macrophages, including EsxB-specific clonotypes that may represent especially relevant targets for pathogen clearance. Notably, TCRs from distinct EsxB-specific GLIPH2 groups (R%SGGEAK%NI and RA%GGEAKNI vs. RKQG%E) differed in their ability to recognize bystander macrophages. Interestingly, we previously identified RKQG%E TCRs in response to peptide-treated but not infected macrophages^35^. These data indicate that that recognition may depend more on the specific TCR-epitope pair than on the abundance of transferred antigen. Additional TCR sequencing may reveal other EsxB-specific and Mtb-specific clonotypes among responses to bystander macrophages.

In summary, our study demonstrates that human memory CD4⁺ T cells can recognize non-infected bystander macrophages through presentation of soluble Mtb antigens transferred from infected cells. Although antigen-specific, many of these T cell responses show attenuated effector cytokine production, suggesting that decoyed T cells could be sidelined from protective immunity. Using scRNAseq, we identified a distinct repertoire of Mtb-specific human memory CD4⁺ T cell clonotypes that recognize non-infected macrophage bystanders, even when separated by several millimeters from infected cells in our transwell system. Identifying antigens and peptides preferentially or exclusively presented by infected macrophages, and defining the T cell phenotypes they elicit, may inform TB vaccine design^60^. Defining how antigen transfer shapes T cell responses within human granulomas will be critical for identifying correlates of protective immunity and for guiding rational vaccine and immunotherapeutic strategies against TB.

## Online Methods

### Human participants

Ten healthy individuals, ages 23–69 years, who self-identified as having latent Mtb infection (LTBI) volunteered for this study. Participants represented African, Asian, White, and Latinx ethnicities. LTBI status was initially determined by a positive tuberculin skin test (TST) of at least 10 mm, a positive IFNγ release assay (IGRA), or both, and was verified using QuantiFERON-TB Gold Plus (Qiagen, Hilden, Germany). Peripheral blood was collected only after written informed consent was obtained and the risks and benefits of study participation were explained. De-identified sample IDs were used for all samples, and no personally identifying information was used for this study. No participants had a history of active TB disease or symptoms suggestive of current disease, including cough, night sweats, fever, or weight loss. Seven participants had received BCG vaccination during infancy (>20 years before participation). All protocols involving human subjects were approved by the Institutional Review Board of University Hospitals Cleveland Medical Center, and informed consent was obtained from all participants. Ten additional HLA-typed leukapheresis products (leukopaks) from healthy male and female volunteers, ages 23–53 years, were purchased from AllCells (Alameda, CA, USA) for experiments performed without T cells or requiring HLA-matched macrophages for TCR-transduced SKW-3 cells. These participants were distinct from the 10 individuals with LTBI described above. No influence of sex on the results was observed, although sample sizes were limited.

### Bacterial culture and infection

Aliquots of H37Rv (NR-13648, BEI Resources) and YFP-Rv (generated by the laboratory of Dr. Christopher M. Sassetti, UMass Chan Medical School, Worcester, MA, USA) were maintained for 6–7 d in Difco Middlebrook 7H9 broth (BD Diagnostics) supplemented with 10% OADC (Millipore), 0.2% glycerol, and 0.05% Tween-80. YFP-Rv cultures were additionally supplemented with hygromycin C (Sigma). Mid-log-phase bacterial cultures (OD_600_ 0.3–0.8) were used to prepare the inoculum after filtration through a 0.5 μm strainer (Millipore). Macrophages were inoculated at the indicated multiplicity of infection (MOI, 0.5–5) for 4 h, followed by three washes with 1X DPBS (Corning) to remove extracellular mycobacteria. For transwell experiments, upper chambers were added to each well only after inoculated cells had been washed three times. Infected cells, including those exposed to bystander cells, were incubated in pre-warmed cRPMI medium [RPMI 1640 supplemented with 10% heat-inactivated fetal bovine serum, 1 mM L-glutamine, 1% sodium pyruvate, 1% non-essential amino acids, 1% HEPES buffer, 0.1% β-mercaptoethanol, and 0.1% 2 M NaOH (Gibco)] at 37°C and 5% CO_2_ for 20–24 h before co-culture with CD4^+^ T cells.

For CFU assessment, Mtb-infected or bystander macrophages, including flow-sorted YFP^-^macrophages, were lysed for 5 min in 1% Triton X-100 in 1X DPBS at 24 h post-infection or immediately after sorting. Lysates were serially diluted in 0.02% Tween-80 in 1X DPBS, and 100 μL aliquots of each dilution were spread onto Difco Middlebrook 7H10 agar plates (BD Diagnostics). Plates were incubated for 18–22 d at 37°C before CFU enumeration. All experimental procedures using virulent Mtb strains were conducted under Biosafety Level 3 (BSL-3) conditions using techniques and protocols approved by the Institutional Biosafety Committee and BSL-3 Advisory Committee of Case Western Reserve University.

### Generation of human monocyte-derived macrophages and memory CD4^+^ T cell isolation

PBMCs were collected from healthy individuals with LTBI from the Cleveland, Ohio, USA area who had no history of active tuberculosis. LTBI status was determined by positive TST and/or IGRA results performed at Case Western Reserve University. CD14^+^ cells were isolated from PBMCs by positive immunomagnetic selection (Miltenyi Biotec) and used for macrophage differentiation, while the CD14^-^ fraction was cryopreserved at -80°C. MDMs were differentiated for 6 d with GM-CSF or M-CSF (PeproTech) to generate M1- or M2-like macrophage phenotypes, respectively^31^. On day 3, half of the medium was replaced with fresh medium containing the same cytokine to maintain differentiation. Autologous memory CD4^+^ T cells were isolated from thawed CD14^-^ fractions from the corresponding donors after resting for at least 3–4 h, or overnight, using a Human Memory CD4 T Cell Isolation Kit (Miltenyi Biotec) based on negative immunomagnetic selection, according to the manufacturer’s instructions. Isolated cells were subsequently used for downstream experiments.

### Flow sorting of YFP-Rv infected macrophages

To flow sort Mtb-infected macrophages, M1-like MDMs were infected with YFP-Rv at an MOI of 1 for 24 h. On the day of sorting, supernatants were collected, and infected cells were washed once with 1X DPBS (∼1 mL/well in 24-well plates), followed by incubation with 250 μL/well Accutase (Gibco) at 37°C and 5% CO_2_ for 10 min. Accutase was neutralized with 250 μL cold cRPMI, and cells were resuspended. Each well was then washed with 500 μL cRPMI to maximize cell recovery. Macrophages were pooled into 15 mL conical tubes, centrifuged at 350 x g for 10 min at 4°C, and stained with Human TruStain FcX (BioLegend) and BV785-conjugated anti-CD11b antibody (clone ICRF44; BioLegend) for 20 min at 4°C in the dark. Cells were washed with AutoMACS Running Buffer (Miltenyi Biotec), resuspended at approximately 2 x 10^6^ cells per 200 μL of the same buffer, and filtered through the strainer of a 40 μm filter-top tube (Corning). The strainer was washed with 200 μL AutoMACS Running Buffer to maximize recovery. Approximately 2 min before sorting, 7-AAD dye (BioLegend) was added to each sample at a 1:100 dilution to exclude dead cells (7-AAD^+^). Sorted cells were collected into polypropylene tubes (Corning) containing 1 mL RPMI supplemented with 20% fetal bovine serum. Macrophage sorting was performed on a Sony MA900 Cell Sorter (Sony Biotechnology) using a 130 μm nozzle chip according to the manufacturer’s instructions. Non-infected M1-like macrophages were used for single-color compensation controls, and YFP-Rv-infected macrophages were used for YFP single-color compensation. Macrophages were sorted in purity mode. Collection tubes containing YFP^-^ and YFP^+^ macrophage fractions were centrifuged at 350 x g for 10 min at 4°C, and cells were resuspended in pre-warmed cRPMI and plated at 50,000 macrophages/well in 96-well flat-bottom plates. Approximately 100,000–200,000 autologous memory CD4^+^ T cells were then added, along with ultra-low endotoxin azide-free (Ultra-LEAF) anti-CD40 mAb (clone W17212H; BioLegend, San Diego, CA, USA). Non-infected macrophages harvested using Accutase, washed, and replated in 96-well flat-bottom plates, as well as a cocktail of anti-MHC-II blocking mAbs (see below), were used as negative controls for T cell activation.

### T cell-macrophage co-culture assays

Isolated memory CD4^+^ T cells were resuspended in pre-warmed cRPMI. For transwell co-cultures, 200,000–250,000 T cells were added to bystander macrophages in upper chambers containing 6.5-mm transwell supports with 0.4-μm tissue culture-treated polycarbonate membranes (Corning Costar, Cat. No. 3413; Corning Inc., Corning, NY, USA) and 50,000 macrophages/insert in 24-well plates. Alternatively, T cells were added to flow-sorted YFP^-^ or YFP^+^ macrophages replated immediately after sorting in 96-well flat-bottom plates (50,000 macrophages/well), or directly to Mtb-infected macrophages in 24-well plates (250,000 macrophages/well). T cell-macrophage co-cultures were incubated at 37°C and 5% CO_2_ for 16– 18 h before harvest for flow cytometry. To facilitate detection of CD40L expression on CD4^+^ T cells, Ultra-LEAF anti-CD40 blocking mAb (clone W17212H; BioLegend) was added at a final concentration of 0.5 μg/mL throughout the co-culture period. To control for cytokine-mediated CD4^+^ T cell activation, a cocktail of two anti-MHC-II blocking mAbs, Ultra-LEAF anti-HLA-DR (clone L234) and Ultra-LEAF anti-HLA-DR/DP/DQ (clone Tü39; BioLegend), was added to control wells, each at a final concentration of 25 μg/mL throughout the co-culture period. AIM^+^ memory CD4^+^ T cells were enumerated by flow cytometry after gating on live (Live/Dead^Lo^) CD3^+^ CD4^+^ CD45RA^Lo^ cells that co-expressed CD69 and CD40L.

To assess T cell responses to conditioned supernatant-treated macrophages, non-infected M1-like macrophages were treated for 24 h before co-culture with autologous memory CD4^+^ T cells using one of the following fractions: 0.22 μm-filtered intact supernatants from infected macrophages, exosomes isolated using the EV Isolation Kit Pan, human (Miltenyi Biotec), exosome-depleted fractions (see below), or exosome-BV-depleted conditioned supernatants (see below). A total of 100,000–200,000 T cells were co-incubated with macrophages treated with each supernatant fraction for 16–18 h at 37°C and 5% CO_2_. BD FastImmune Co-Stimulatory Antibodies [anti-CD28 (clone L293) and anti-CD49d (clone L25); Waters Biosciences] were added according to the manufacturer’s instructions, together with Ultra-LEAF anti-CD40 mAb blockade with or without the anti-MHC-II mAb blocking cocktail throughout the co-culture period.

### Immunophenotyping, isolation, and depletion of Extracellular Vesicles (EVs)

TexMACS Medium (Miltenyi Biotec) was used to culture non-infected or Mtb-infected M1- and M2-like macrophages (H37Rv at MOI 4) to prevent contamination with fetal bovine serum-derived exosomes. Samples were frozen at -80°C until further use. Immunophenotyping of human exosomes was performed using the MACSPlex EV Kit (Miltenyi Biotec) according to the manufacturer’s instructions. Human exosomes were isolated from conditioned media collected from M1-like macrophages 48 h post-infection (H37Rv at MOI 4) using the EV Isolation Kit Pan, human (Miltenyi Biotec), based on positive immunomagnetic selection according to the manufacturer’s protocol. Isolated exosomes were eluted in cRPMI and resuspended to the original volume of the starting supernatant to approximate the physiological exosome density present during infection *in vitro*.

For EV depletion, conditioned supernatants were pre-cleared according to the EV Isolation Kit Pan, human (Miltenyi Biotec) protocol. Samples were filtered through 0.22 μm Spin-X tubes (Corning) and passed through immunomagnetic columns for exosome isolation, and exosome-depleted flow-through conditioned media were collected. Exosome depletion was confirmed using the MACSPlex EV Kit (Miltenyi Biotec). To further deplete remaining EVs, including bacterial vesicles (BVs), exosome-depleted flow-through fractions were ultracentrifuged at 100,000 x g for 120 min at 10°C using a Type 50.2 Ti fixed-angle rotor in an Optima^TM^ XE-90 preparative ultracentrifuge (Beckman Coulter Life Sciences, Brea, CA, USA). The top 75% of the supernatant was collected and designated exosome-BV-depleted, while the remaining 25% was discarded to avoid pellet disruption.

### Flow cytometric analysis

Immunostaining and sample preparation for sorting were performed as described previously^30,35^. Stained samples were subsequently fixed with 1% paraformaldehyde (PFA) for 1 h prior to removal from the BSL-3, in accordance with our biosafety-approved protocol. Fixed cells were then washed and resuspended in AutoMACS Running Buffer (Miltenyi Biotec) for analysis on a BD LSRFortessa X-20 Cell Analyzer (BD, Franklin Lakes, NJ, USA). For MDMs, YFP fluorescence and expression of individual surface-markers were quantified after gating on LiveDead^Lo^ CD11b^+^ single cells. For CD4^+^ T cell analyses, the proportion of cells co-expressing CD69 and CD40L was determined after gating on LiveDead^Lo^ CD4^+^ CD45RA^Lo^ single cells. The proportion of activated T cells expressing GM-CSF, IFNγ, IL-2, or TNF was determined after gating on LiveDead^Lo^ CD4^+^ CD45RA^Lo^ CD69^+^ CD40L^+^ single cells.

Multicolor flow panels for T cells included the following fluorochrome-conjugated antibodies: anti-human CCR6-BV421 (clone G034E3), CCR4-BV605 (L291H4), CD45RA-BV650 (HI100), CD4-BV786 (RPA-T4), CXCR3-PerCP-Cy5.5 (G025H7), CD40L (CD154)-PE-Dazzle594 (24-31), 41BB (CD137)-PECy7 (4B4-1), HLA-DR-APC-Fire750 (I.243), CD45RO-AF700 (UCHL1), and anti-mouse TCRβC-PE (H57-597) (for SKW-3 cell assays) from Biolegend. Anti-human CD25-BB515 (M-A251), CD69-BUV396 (FN50), and CD3-BUV737 (UCHT1) from Waters (formerly BD) Biosciences. Anti-human TNF secretion assay-PE, anti-IFNγ secretion assay-APC, anti-GM-CSF secretion assay-PE, and anti-IL-2 secretion assay-APC from Miltenyi Biorec. Live/Dead Fix Aqua (Life Technologies).

### Flow sorting of AIM+ T cells

For isolation of AIM+ T cells, memory CD4^+^ T cells were harvested after co-culture with Mtb-infected or bystander macrophages for 16-18 h in transwell plates. Immunostaining and sample preparation for sorting were performed as described^30^. The Sony MA900 Cell Sorter (Sony Biotechnology) was configured according to manufacturer’s instructions. T lymphocytes were sorted from Live CD4^+^ CD45RA^Lo^ based on co-expression of CD69 and CD40L in “purity” mode. To facilitate detection of CD40L expression on CD4^+^ T cells, Ultra-LEAF anti-CD40 blocking antibody was added at a final concentration of 0.5 μg/mL throughout the T cell-macrophage co-culture period. Flow-sorted memory CD69^+^ CD40L^+^ CD4^+^ T cells were then processed for scRNAseq.

The following fluorochrome-conjugated antibodies were used for flow sorting of AIM+ T cells: Anti-human CD69-BV421 (FN50), CD137-APC (4B4-1) from Waters Biosciences, and CD4-FITC (RPA-T4), CD25-PE (BC96), CD40L-PE-Dazzle-594 (24-31), CD8-BV605 (RPA-T8), CD274-BV711 (29E.2A3), CD45RA-APC-Fire750 (HI100), and 7-AAD viability Staining Solution from Biolegend.

### Sample processing for scRNAseq

The 10X Genomics Chromium Next GEM Single Cell 5’ V2 platform (10X Genomics) was used for sequencing of mRNA from freshly sorted memory CD69^+^ CD40L^+^ CD4^+^ T cells. Sample processing and quality control were performed as described previously^35^, and according to 10X Genomics User Guide. Briefly, activated CD4^+^ T cells were resuspended in 0.04% BSA in PBS and loaded onto the 10X Chromium Controller. RNA was reverse-transcribed, and cDNA was isolated and PCR-amplified according to the 10X Genomics NextGEM 5’ V2 scRNAseq User Guide (Rev D). Quality control and quantification of cDNA and final libraries were performed on either the Agilent fragment analyzer or Agilent 2100 Bioanalyzer (Agilent Technologies, Santa Clara, CA, USA). V(D)J amplification and library preparation, and gene expression library construction were performed using 10X Genomics kits according to the User Guide. Paired-end sequencing was performed by the Cleveland Clinic Lerner Research Institute Genomics Core (Cleveland, OH, USA) using an Illumina NovaSeq 6000 sequencer (Illumina, San Diego, CA, USA) according to the 10X Genomics User Guide. Demultiplexed FASTQ files for TCR sequencing and single-cell gene expression were mapped to the human genome using Cell Ranger Multi V9 (10X Genomics) on the 10X Cloud server and the GRCh38.p14 reference genome.

### Single-cell data analysis

The TCR clonotype analysis used the filtered_contig_annotations.csv output from Cell Ranger containing productive, barcoded, full-length TCR sequences for each sample. The filtered feature matrix containing the gene expression was imported into R v4.5.2 with Seurat v5.3.1^62,63^ and BPCells v0.3.1^64^. The RNA and TCR information was combined using scRepertoire v2.7.2^65^. TCR clonotypes based on CDR3α/β chains were classified as expanded when present, based on 10X cell barcode, in 2 or more cells. Cells with <200 features, >5000 features, >25000 feature counts, or >10% mitochondrial counts were excluded from downstream analyses. For feature normalization, SCTransform^66^ was used to fit model parameters and sctransform_pearson^67^ from BPCells was used to fit the model to the on-disk matrix. The data was scaled by the difference between the cell cycle S score and G2M score using Seurat’s CellCycleScoring function. TCR, long non-coding RNA molecules, and ribosomal subunit genes were removed, if present. Prior to calculating a PCA, variable features matching the regular expression “^TR[ABDG][VDJ]” (all TCR genes) were removed from the variable feature list to reduce the impact of TCR genes on downstream clustering and dimension reduction. Harmony^68^ was used to integrate the samples, and the UMAP embedding and the shared-nearest neighbor graph were calculated using the first 30 harmony dimensions. Louvain clusters were calculated with a resolution of 0.1. MAST v1.36.0^69^ was used for differential expression (Extended Data 4) with the default parameters to FindMarkers. Plots were created using EnhancedVolcano, SCpubr, and ComplexHeatmap^70,71^.

### TCR Sequencing Analysis

A combined list of the unique (and total) TCR clonotypes identified in response to Mtb-infected or bystander macrophages was generated from all experiments which evaluated a response pair. This list was then combined with data from the same participants and other data from Cleveland and South African participants we recently published^35^. A list of all HLA-II alleles for each participant, and a combined list of all CDR3β, CDR3α, Vβ, and Jβ genes, and number of clones, for all TCRs from each participant and experimental condition was generated. For peptide stimulations, only expanded TCRs (≥2 copies) were included. In cases where two CDR3β sequences were identified in a cell, they were separated and each linked to the same CDR3α, V, and J genes for compatibility with GLIPH2. GLIPH2 analysis on the combined list of TCRs (infected and bystander macrophage pairs) was performed as described^36^ using BLOSUM62 restriction for interchangeable amino acids. GLIPH2 groups considered sufficiently robust for further analysis were selected based on the following criteria: Representation from ≥3 individuals and ≥ 2 unique CDR3β sequences within a group, TCR Vβ gene homology score (p < 0.05), CDR3β length distribution score (p < 0.01), clonal expansion score (p < 1), HLA association score (p < 0.05), and Fisher exact score (p < 1) for distinct CDR3β motifs as reported previously^35,41^. Using the jvenn Venn diagram builder^72^, GLIPH2 groups containing TCRs linked to a response to infected macrophages, bystander macrophages, or only peptides or lysate were compared. TCR sequences within GLIPH2 groups were cross-referenced with TCRs annotated in the immune epitope database (IEDB) using TCR Match^51^, and we removed entire GLIPH2 groups that contained any CDR3βs that contained ≥ 97% homology with TCRs previously annotated as specific for viral antigens in other studies^35^ (Extended Data 1). Stacked bar plots were generated in Prism to express the number of copies of each unique TCR within a GLIPH2 group sequenced from the responses to infected or bystander macrophages from each of the 5 participants.

### SKW-3 cell-macrophage co-culture assays

SKW-3 cells (Cytion, Heidelberg, Germany; Cat# 300343) bearing human Mtb-specific TCRs were generated previously by lentiviral transduction using representative TCRs synthesized from GLIPH2 groups linked to responses to Mtb peptides or infected macrophages^35^. Approximately 200,000 TCR-transduced SKW-3 cells were added to bystander macrophages in the upper chambers of 24-well transwell plates (50,000 macrophages/insert), or directly to Mtb-infected macrophages and control conditions (50,000 macrophages/well in 96-well flat bottom plates). SKW-3 cell-macrophage co-cultures were incubated at 37°C and 5% CO_2_ for 16-18 h prior to harvest for flow cytometry. SKW-3 cells express CD69 upon TCR engagement, used to detect activation. To control for cytokine-mediated CD69 upregulation, the anti-MHC-II blocking mAb cocktail was again added at a final concentration of 25 μg/mL each, throughout the co-culture period.

### TCR epitope identification using the IMPAc-TB peptide library

The antigen specificity of a representative TCR from the SA%RDQP GLIPH2 group, described previously^35^, was identified screening TCR-transduced SKW-3 cells against autologous EBV-transformed B cells loaded with pools of peptides. Although this TCR did not respond to the MTB300 peptide megapool, an IMPAc-TB library of overlapping peptides from secreted Mtb antigens was generated. The IMAPc-TB peptide library is comprised of 68 proteins^73^ arrayed as 15mers overlapping by 11 amino acids. The sequences are based on H37Rv. To ensure that we accounted for sequence variation across as many known Mtb strains as possible, we analyzed each 15mer for variance among approximately 50,000 strains of Mtb and 15mers and sequence variants were included in the library. A total of 8024 15mers were synthesized, split into 127 pools of between 50 and 100 peptides (Extended Data 2). For initial TCR screening, small aliquots of these 127 pools were combined into 26 super-pools containing ∼300 peptides each. A positive hit was deconvoluted into the original individual pools, followed by new synthesis of individual crude peptides spanning the amino acid sequences of the Mce3A antigen likely to contain the epitope.

### Mass spectrometry analysis of conditioned supernatants

Mtb-infected (H37-Rv at MOI 4) M1- or M2-like MDMs cultured in TexMACS GMP Medium (Miltenyi Biotec) without phenol red were incubated for 48 h at 37°C and 5% CO_2._ Supernatants were collected, filtered through a 0.22 μm strainer, and stored at -80°C until further use. Samples were submitted to the Proteomics Core of Cleveland Clinic Foundation for mass spectrometry analysis. Samples were thawed at room temperature, and two 100 μL aliquots of each sample were removed for processing by standard in-solution digestion or by digestion using the S-Trap protocol.

For the in-solution digestion, the samples were diluted with 100 μL Tris HCl/Urea buffer, reduced with DTT, and alkylated with iodoacetamide prior to precipitation by the addition of 1mL cold acetone overnight. The proteins were digested using 100 μL trypsin in 50 mM TEAB and incubated overnight at room temperature. The digestion was stopped using 5 μL of 20% TFA, dried in a Speedvac and reconstituted in 30 μL 0.1% FA. Prior to LC-MS, the samples were passed through a 0.22 μm filter.

For the S-Trap digestion, the samples were diluted with 100 μl of S-Trap lysis buffer (10% SDS in 100mM TEAB). These samples were sonicated, reduced with DTT, and alkylated with iodoacetamide prior to acidification with phosphoric acid and dilution with 100 mM TEAB in 90% methanol. The proteins were loaded onto S-Trap tubes and centrifuged, then washed with 100 mM TEAB in 90% methanol. After washing, the proteins were digested by the addition of trypsin at a 1:25 ratio and incubated overnight at room temperature. Peptides were eluted off the S-trap columns using 50 mM TEAB, 0.2% formic acid, and 50% acetonitrile. The digestion was stopped by the addition of 5 μL of 20% TFA. The elutions were combined and dried in a Speedvac and reconstituted in 30 μL 0.1% FA. Prior to LC-MS, the samples were passed through a 0.22 μm filter.

The Bruker TimsTof Pro2 Q-Tof liquid chromatography mass spectrometry (LC-MS) system operating in positive ion mode, coupled with a CaptiveSpray ion source (Bruker Daltonik GmbH, Bremen) uses a Bruker 15 cm x 75 μm id C18 ReproSil AQ, 1.9 μm, 120 Å reversed-phase capillary HPLC a column. 1 μL volumes of the extract were injected and the peptides eluted from the column by an acetonitrile/0.1% formic acid gradient at a flow rate of 0.3 μL/min were introduced into the source of the mass spectrometer. The Parallel Accumulation–Serial Fragmentation DDA method was used to select precursor ions for fragmentation with a TIMS-MS scan followed by 10 PASEF MS/MS scans. The TIMS-MS survey scan was acquired between 0.60 and 1.6 Vs/cm^2^ and 100–1,700 m/z with a ramp time of 166 ms. The total cycle time for the PASEF scans was 1.2 s and the MS/MS was performed with a collision energy between 20 eV (0.6 Vs/cm2) and 59 eV (1.6 Vs/cm2). Precursors with 2–5 charges were selected with the target value set to 20,000 AU and intensity threshold to 2,500 AU. Precursors were dynamically excluded for 0.4 s. Mtb and human proteins were identified by matching experimentally observed peptide spectra against reference protein sequence databases, inferring protein identities when ≥ 2 peptides from the same protein were identified (Extended Data 3).

### APC light fixation assays

M1-like macrophages were differentiated for 6 d in 96-well flat-bottom plates (50,000 cells/well) and remained uninfected before fixation. Cells were treated with 100 μL/well of 0.25% paraformaldehyde (PFA) diluted in warm RPMI for 15 min at 37°C and 5% CO_2_. PFA was removed, and fixed cells were washed three times with 200 μL/well of warm RPMI before addition of 200 μL/well 0.2 M lysine diluted in warm RPMI for 15 min at 37°C and 5% CO_2_ to quench residual reactive aldehyde groups. Treated cells were washed three times with warm cRPMI before exposure to intact 0.22 μm-filtered conditioned supernatants. Intact supernatants were harvested from cultures of M1-like macrophages infected at an MOI of 3 for 48 h, filtered through a 0.22 μm strainer, and added to MDMs with or without prior PFA fixation for 1 or 16 h before co-culture with TCR-transduced cell lines. In parallel, cognate peptide (EspA_301-315_) or whole Mtb cell lysate (1 μg/mL) was added to MDMs with or without prior PFA fixation for 1 or 16 h, respectively, before co-culture with TCR-transduced cell lines. After treatment with intact supernatants or exogenous Mtb antigens, cells were washed with 1X DPBS, and each TCR-transduced cell line was added and incubated for 24 h at 37°C and 5% CO_2_.

### Quantification and statistical analysis

Sample sizes for *ex vivo* T cell activation co-culture experiments were based on feasibility of obtaining sufficient numbers of monocytes and autologous primary human memory CD4^+^ T cells from individual participants with LTBI, while prioritizing paired within-donor comparisons across experimental conditions. All statistical comparisons were made using 2–4 technical replicates per condition to ensure reproducible detection of donor-matched differences in T cell activation. Flow cytometry data were analyzed using FlowJo v10 (BD Biosciences, San Diego, CA, USA). For each sample, the proportion of cells expressing each marker (and median fluorescence intensity (MFI) was quantified and compared between groups. Statistical comparisons were performed using raw values or absolute differences in MFI between conditions, as indicated. Data were tested for normality using the Shapiro-Wilk or Kolmogorov-Smirnov tests in Prism V11 (GraphPad, San Diego, CA, USA). If normally distributed, a One-way ANOVA with Šidák’s post-test and correction for multiple comparisons was used to determine statistical significance when comparing 3 or more groups, where indicated. A Kruskal–Wallis test with Dunn’s post-test was used when non-parametric data were analyzed. Fisher Exact score, CDR3 length distribution score, Vβ gene homology score, clonal expansion score, and HLA association score for each GLIPH2 similarity group were generated by GLIPH2 and used to select robust groups, as described^36,41^. For single-cell transcriptomics data, MAST (adjusted p-value < 0.05 and abs(log_2_ fold change) > 0.5)) was used to determine differentially expressed genes.

## Acknowledgments

The following reagents were obtained through BEI Resources, NIAID, NIH: *Mycobacterium tuberculosis*, Strain H37Rv, whole cell lysate, NR-14822 and *Mycobacterium tuberculosis* strain H37Rv, NR-13648. This work was supported by National Institutes of Health (NIH) grants K08 AI163407 and R01 AI187662 (to S.M.C.), R21 AI167675 (to S.M.C and M.L.F), and research funds from University Hospital Cleveland Medical Center (to S.M.C.). Intellectual support was provided by NIH IMPAc-TB Center contract 75N93019C00071.The IMPAc-TB peptide library was constructed with support from NIH IMPAc-TB Center contract 75N93019C00070 (to D.M.L. and D.A.L.). BSL-3 core facilities were supported by the Case Western Reserve University School of Medicine. We are grateful to Dr. Brian Cobb for thoughtful scientific discussion, to Sophia Onwuzulike for assistance with BSL-3 cell sorting, and to Belinda Willard of the Cleveland Clinic Foundation Proteomics and Metabolomics Core for assistance with mass spectrometry. Graphical abstract and select figure panels were created with Biorender.com.

## Author contributions

Conceptualization, V.G.S., S.M.C.; Methodology, V.G.S., M.L.F., C.V.H., S.M.C.; Investigation, V.G.S., R.S., D.P.G., S.M.R., S.M.C.; Resources, V.G.S., S.M.R., D.P.G., G.S., D.M.L., D.A.L., S.M.B., C.M.S., B.L., B.D.B., W.H.B., C.V.H., S.M.C.; Formal Analysis, V.G.S., R.S., S.M.R., S.M.C.; Writing-original draft preparation, V.G.S., R.S., and S.M.C.; Writing-review and editing, V.G.S., R.S., S.M.R., D.P.G., G.S., D.M.L., D.A.L., C.M.S., B.D.B., M.L.F., W.H.B., C.V.H., S.M.C.; Visualization, V.G.S., R.S., S.M.R., and S.M.C.; Supervision, S.M.C.; Funding Acquisition, M.L.F., D.M.L., D.A.L., and S.M.C.

## Data Availability

Raw single-cell sequencing data used in Fig. 3, 5, and 6 will be publicly available upon publication through NCBI Gene Expression Omnibus (GEO) accession number <u>GSE341732</u>. GLIPH2 outputs of TCR sequencing data used for Fig. 3 are available in Extended Data 1. The sequences of all peptides in the IMPAc-TB peptide library are available in Extended Data 2. Results from mass spectrometry of conditioned supernatants are available in Extended Data 3. Genes differentially expressed by each Louvain cluster in Fig. 5 are available in Extended Data 4. Other data used to generate figures will be publicly available upon publication through figshare at https://doi.org/10.6084/m9.figshare.33868102. Any additional information required to reanalyze the data reported in this paper is available from the corresponding author upon reasonable request. This paper does not report original code.

## Supplementary Figure Legends

**Supplementary Figure 1 (related to Fig. 1, 3 and 6). Gating strategy for YFP-based macrophage sorting assay and AIM+ CD4^+^ T cells. (A)** Representative flow cytometry plots of LiveDead^Lo^ CD11b^+^ YFP^+^ M1-like macrophages 24 h post infection with YFP-Rv (MOI 1). (**B)** Representative flow cytometry plots of the gating strategy used to identify memory CD4^+^ T cells activated 16-18 h after co-culture with Mtb-infected or bystander M1-like macrophages.

**Supplementary Figure 2 (related to Fig. 3 and 4). TCR specificity identification of Rv1966_111-125_-specific cell line and recognition of BYST macrophages in transwell system. (A)** Table containing each TCR cloned into SKW-3 cells used in experiments. **(B, C)** Summary bar graphs of the proportion of SKW-3 cells transduced with a TCR from the SA%RDQP GLIPH2 group expressing CD69 (gated on LiveDead^Lo^ CD4^+^ TCRβ^+^), 16-18 h after co-culture with autologous EBV-transformed B cells loaded with **(B)** 26 super-pools of IMPAcTB peptide pools containing overlapping peptides from 68 Mtb antigens (Extended Data 2), and **(C)** 20 overlapping peptides of Rv1966 from super-pool 11. **(D)** Representative summary bar graph of EspC_74-88_-specific TCR-transduced SKW-3 cells expressing CD69 16-18 h after co-culture with INF or Non-Inf macrophages, or in upper transwell chambers with BYST macrophages exposed to infected macrophages for 0 h, 24 h and 48 h after the Mtb inoculum was washed out of lower chambers. Data represent single replicates from 2 independent experiments.

**Supplementary Figure 3 (related to Fig. 4). Gating strategy for immunophenotyping of human exosomes by flow cytometry-based assay.** Representative flow cytometry plots of the gating strategy for identification of CD9^+^, CD63^+^, CD81^+^, and HLA-DR/DP/DQ^+^ exosomes in 0.22 μm-filtered supernatants conditioned by infected M1- or M2-like MDMs by flow cytometry.

**Supplementary Figure 4 (related to Fig. 5). Genes differentially expressed by memory CD4^+^ T cells responding to INF or BYST macrophages in Louvain clusters.** Heatmap representing log-normalized counts of differentially expressed genes of the flow-sorted activated CD4^+^ T cell populations in response to INF or BYST macrophages after quality control and integration. Data are combined from 2 experiments containing samples from 5 individuals.

## Supplementary Data Files

**Extended Data 1** (related to Fig. 3) lists the output of GLIPH2 clustering of TCRs.

**Extended Data 2** (related to Fig. 3) lists the overlapping peptides that comprise the pools representing 68 antigens used to screen TCRs cloned from each GLIPH2 group.

**Extended Data 3** (related to Fig. 4) lists the proteins identified in conditioned supernatants by mass spectrometry.

**Extended Data 4** (related to Fig. 6) lists the DEGs from each Louvain cluster of activated CD4^+^ T cells from single-cell transcriptomics data.

