## Supplementary Figures for "Human memory CD4^+^ T cells recognize non-infected macrophage bystanders exposed to *Mycobacterium tuberculosis*-infected cells"

### Supplementary Figure 1

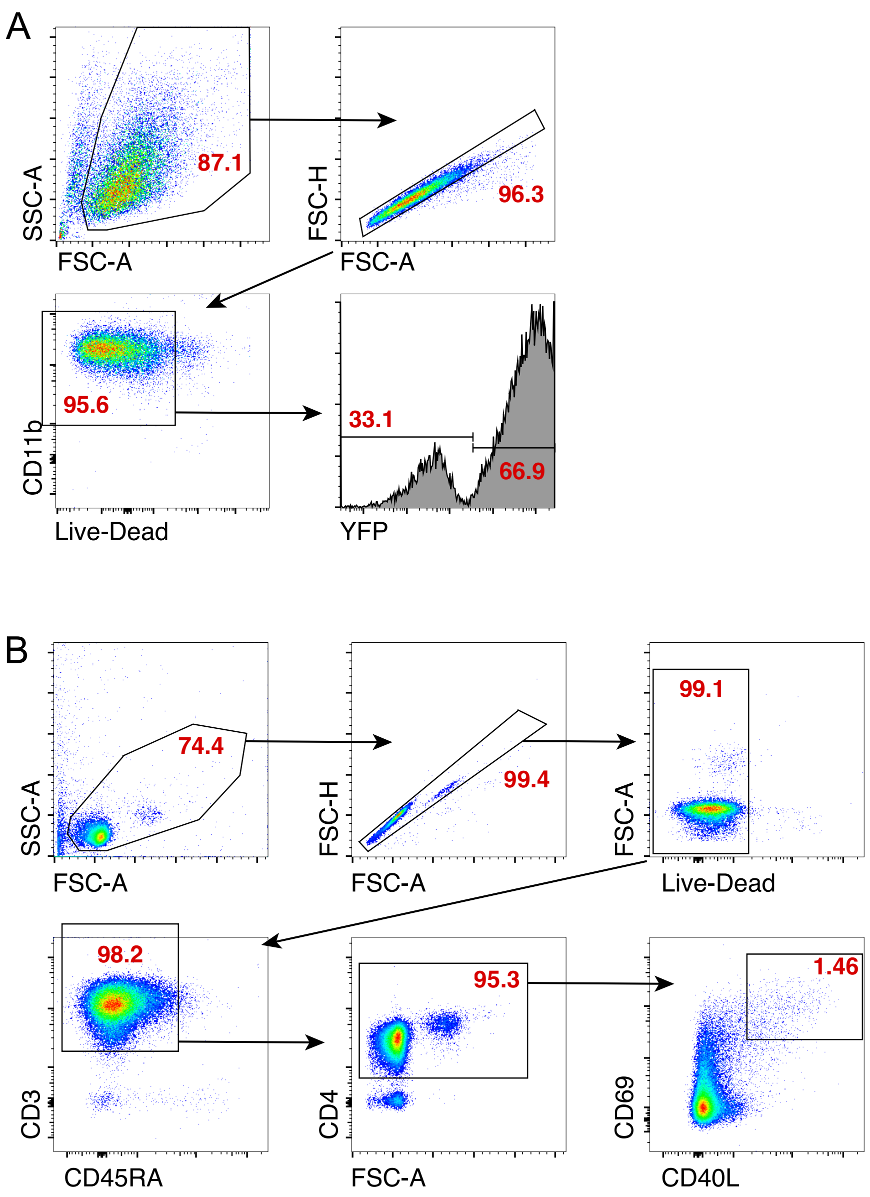


**Supplementary Figure 1** (related to Fig. 1, 3 and 6). Gating strategy for YFP-based macrophage sorting assay and AIM+ CD4+ T cells. (A) Representative flow cytometry plots of LiveDeadLo CD11b+ YFP+ M1-like macrophages 24 h post infection with YFP-Rv (MOI 1). (B) Representative flow cytometry plots of the gating strategy used to identify memory CD4+ T cells activated 16-18 h after co-culture with Mtb-infected or bystander M1-like macrophages.

### Supplementary Figure 2

A

| GLIPH2 motif(s) | CDR3α | CDR3β | HLA restriction | Epitope | Mtb protein |
|---|---|---|---|---|---|
| SLGTESN%P | CALKFSGGYNKLIF | CASSLGTESNQPQHF | DPB1*13:01 | STRQALRPRADGPVG | EspA (Rv3616) |
| SSPGQGG%NYG | CATARTGANNLFF | CASSSPGQGGSNYGYTF | DRB1*07:01 | VDLAKSLRIAAKIYS | EspC (Rv3615c) |
| RKQG%E | CIVRVGQGRAGNNRKLIW | CASRKQGHEQFF | DRB1*11:02 | EISTNIRQAGVQYSRADEEQ | EsxB (CFP-10/Rv3874) |
| SA%RDQP | CAVHGSSNTGKLIF | CASSAHRDQPQHF | DRB4*01:03 | AFGAKFVDLVMPQNP | Mce3A (Rv1966) |

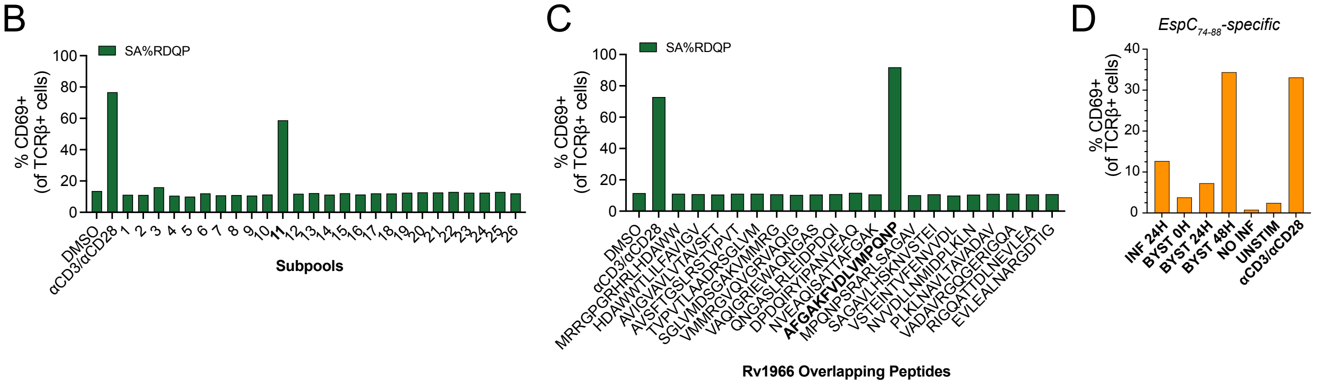


**Supplementary Figure 2** (related to Fig. 3 and 4). TCR specificity identification of Rv1966111-125-specific cell line and recognition of BYST macrophages in transwell system. **(A)** Table containing each TCR cloned into SKW-3 cells used in experiments. (B, C) Summary bar graphs of the proportion of SKW-3 cells transduced with a TCR from the SA%RDQP GLIPH2 group expressing CD69 (gated on LiveDeadLo CD4+ TCR +), 16-18 h after co-culture with autologous EBV-transformed B cells loaded with **(B)** 26 super-pools of IMPAcTB peptide pools containing overlapping peptides from 68 Mtb antigens (Extended Data 2), and **(C)** 20 overlapping peptides of Rv1966 from super-pool 11. **(D)** Representative summary bar graph of EspC74-88-specific TCR-transduced SKW-3 cells expressing CD69 16-18 h after co-culture with INF or Non-Inf macrophages, or in upper transwell chambers with BYST macrophages exposed to infected macrophages for 0 h, 24 h and 48 h after the Mtb inoculum was washed out of lower chambers. Data represent single replicates from 2 independent experiments.

### Supplementary Figure 3

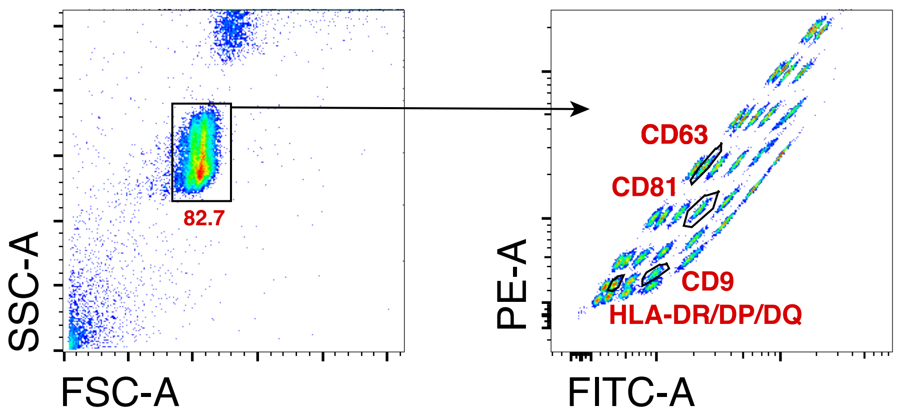


**Supplementary Figure 3** (related to Fig. 4). Gating strategy for immunophenotyping of human exosomes by flow cytometry-based assay. Representative flow cytometry plots of the gating strategy for identification of CD9+, CD63+, CD81+, and HLA-DR/DP/DQ+ exosomes in 0.22 m-filtered supernatants conditioned by infected M1- or M2-like MDMs by flow cytometry.

### Supplementary Figure 4

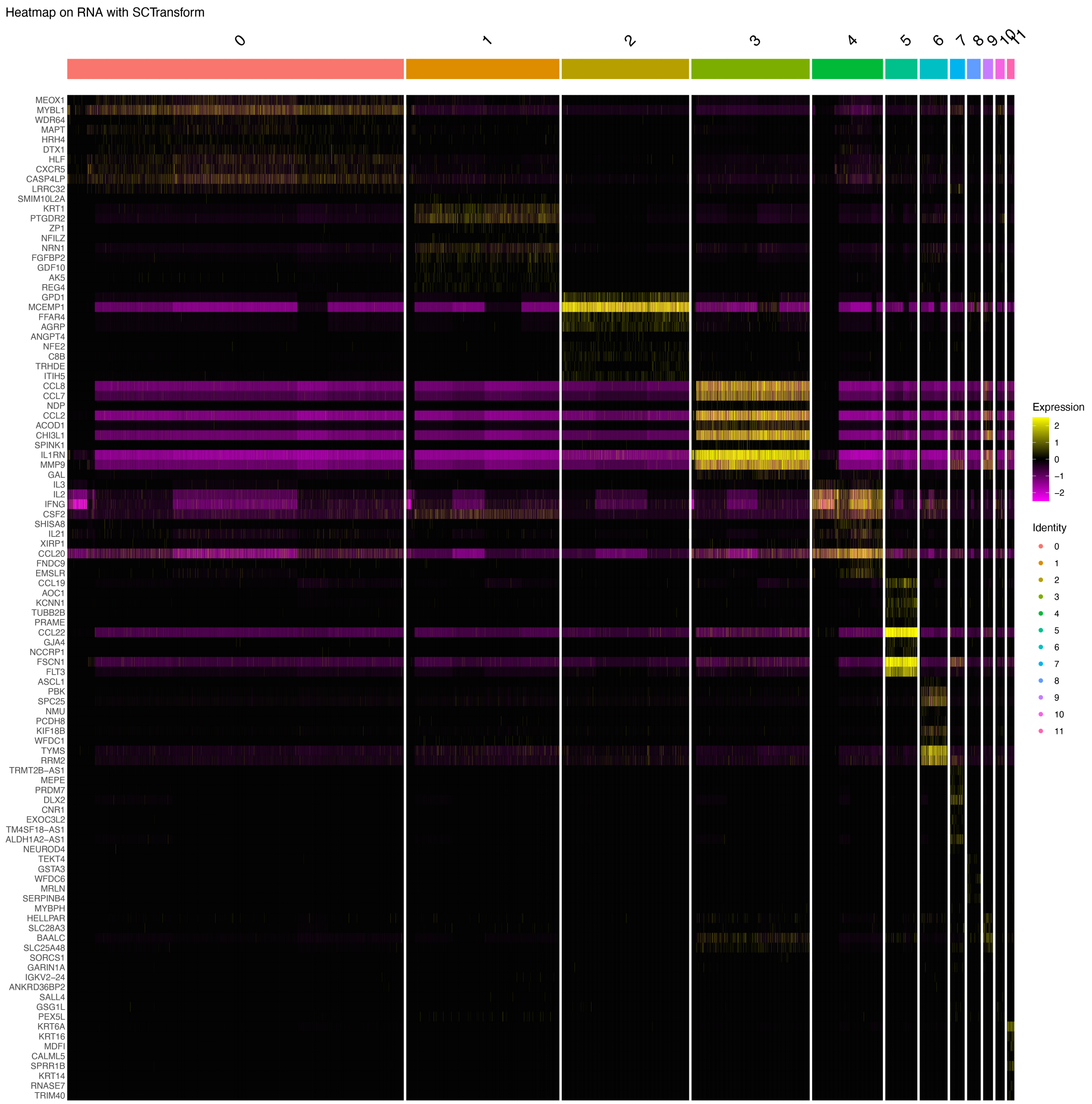


**Supplementary Figure 4** (related to Fig. 5). Genes differentially expressed by memory CD4+ T cells responding to INF or BYST macrophages in Louvain clusters. Heatmap representing log-normalized counts of differentially expressed genes of the flow-sorted activated CD4+ T cell populations in response to INF or BYST macrophages after quality control and integration. Data are combined from 2 experiments containing samples from 5 individuals.
